# Asymmetric photosynthetic responses to hydrothermal variations between the two halves of the year in the Amazon rainforest

**DOI:** 10.1101/2025.09.03.673977

**Authors:** Yuchuan Yang, Yu Shen, Haoxin Tan, Hongjun Yang, Yongchang Ye, Zhanmang Liao, Chaoyang Wu, Josep Peñuelas, Philippe Ciais, Lei Chen

## Abstract

The Amazon rainforest, which stores approximately 120 billion tons of carbon and contributes around 16% of global terrestrial photosynthetic productivity, plays a pivotal role in global carbon cycling. Unlike temperate and boreal forests, tropical forests exhibit a bimodal photosynthetic pattern, characterized by distinct peaks in the first and second halves of the year. However, the intra-annual differences in photosynthetic responses to hydrothermal variations between these two periods in the Amazon rainforest remain largely unexplored. Here, utilizing satellite-derived photosynthetic proxies alongside ground-based flux tower observations from 2001 to 2020, we investigated the differences in photosynthetic responses to hydrothermal variations between the first and second halves of the year in the Amazon rainforest. Our observations revealed weaker temperature limitations but stronger precipitation limitations on photosynthesis in the second half of the year compared to the first half. Temperature constraints on photosynthesis have progressively weakened in both periods, while precipitation limitations have intensified, particularly in the latter half. Although the optimal temperature for photosynthesis is higher in the second half of the year, it is reached earlier, resulting in a sharper decline in photosynthetic productivity over the past two decades. Our findings reveal a shift from temperature to precipitation limitation in the Amazon, underscoring intra-annual asymmetry in vulnerability to intensifying heatwaves and droughts and calling for its explicit integration into management strategies and predictive models.

## Introduction

Photosynthesis in the Amazon rainforest is a key driver of the global carbon cycle, facilitating the uptake of ∼120 billion tons of carbon and contributing to ∼16% of global terrestrial photosynthetic productivity^1,2^. Given this substantial role, even minor fluctuations in Amazonian photosynthetic activity can have far-reaching implications for global carbon dynamics^3^. Over recent decades, warming-induced heatwaves and droughts have triggered widespread tree mortality and forest degradation^4–8^, with some regions shifting from carbon sinks to carbon sources^9–13^. Understanding how photosynthetic activity responds to hydrothermal variations is critical for predicting shifts in Amazonian carbon cycling as well as informing carbon accounting frameworks and land-use policy decisions under future climate scenarios.

Unlike temperate forests, which exhibit a single growing season constrained by cold winters, tropical forests experience year-round favorable conditions^14,15^. Satellite and ground-based observations have revealed that evergreen broadleaved forests in tropical regions, including the Amazon, exhibit a bimodal leaf and photosynthetic phenology, characterized by two distinct peaks in the first and second halves of the year, respectively^14,16–24^. Despite the critical role of hydrothermal conditions in regulating photosynthesis, it remains unclear whether and how the photosynthetic responses to hydrothermal variability differ between these two periods. Additionally, photosynthetic activity often exhibits nonlinear responses to rising temperature. Photosynthesis follows a convex temperature response, peaking at an optimal temperature (TDDD) and declining thereafter^25,26^. Although recent studies suggest that Amazonian forests may already be operating near TDDD^27–29^, it remains uncertain whether TDDD differs between the first and second halves of the year and when temperatures in each period exceed TDDD. Therefore, elucidating the semi-annual cycle of photosynthetic responses to a warming climate is essential for predicting the future trajectory of Amazonian photosynthesis and its impact on the global carbon cycle.

Previous studies assessing the climatic responses of Amazonian photosynthesis have largely relied on the dry-wet seasonal framework^30–34^. However, this classification presents several limitations. First, definitions of dry and wet seasons vary across the Amazon^30,31,35–38^, as some regions lack a distinct dry season according to certain criteria (e.g., monthly precipitation < 100 mm)^35^. Second, the duration of dry and wet seasons fluctuates annually due to climate variability^39–41^, complicating the establishment of a consistent temporal framework. As a result, the traditional dry-wet season division may be inadequate for fully capturing the climatic responses of bimodal photosynthesis in the Amazon rainforest. To address these limitations and provide a more comprehensive framework for analyzing the bimodal photosynthetic pattern, we propose a semi-annual classification based on natural photosynthetic seasonality. This approach offers several advantages: (1) it better aligns with the bimodal phenology of vegetation growth, capturing seasonal variations in photosynthesis more accurately; (2) it further incorporates the integrated effects of climate on photosynthesis, rather than relying solely on hydrological patterns; and (3) it mitigates inconsistencies arising from regional precipitation heterogeneity, providing a spatially consistent framework. Therefore, this semi-annual classification offers a spatially and temporally coherent framework that delineates two complete photosynthetic cycles, facilitates cross-scale comparisons, and captures the coupled hydrothermal influences on photosynthesis.

In this study, we examined intra-annual asymmetry in photosynthetic responses to hydrothermal variability over 2001 – 2020, integrating multiple satellite-derived proxies with ground-based flux tower observations. We first confirmed the existence of a bimodal photosynthetic pattern across the Amazon rainforest. We then calculated and compared the sensitivity of photosynthesis to temperature (S_T_) and precipitation (S_P_) between the first and second halves of the year. We further decoupled the individual effects of temperature and precipitation on S_T_ and S_P_ to clarify the climatic drivers underlying these asymmetric responses. Moreover, we examined the temporal trends of S_T_ and S_P_, identifying historical and projected tipping years for the optimal temperature for photosynthesis (T□□□) in both halves of the year. Finally, we assessed reversals in the temperature effect on interannual photosynthetic variations. We hypothesize that photosynthetic responses to temperature and precipitation differ significantly between the two halves of the year due to distinct hydrothermal conditions. By adopting this semi-annual classification, our study aims to bridge existing knowledge gaps in Amazonian climate-photosynthesis interactions and provide a robust framework for improving carbon cycle modeling in the Amazon rainforest under future climate change.

## Results

### Bimodal photosynthetic pattern across the Amazon rainforest

We first provided robust evidence for the bimodal photosynthetic pattern in the Amazon rainforest using high-temporal-resolution observations from the Advanced Baseline Imager (ABI) onboard the GOES-16 satellite (see Methods). Using a spline-based land surface phenology detection algorithm, we examined the photosynthetic growth patterns and extracted the timing of photosynthetic peaks, represented by the day of year (DOY), from the ABI NBAR EVI2 time series for the Amazon rainforest in 2018 and 2019 (see Methods). We found that bimodal photosynthetic seasonality was prevalent across the Amazon rainforest, with ∼80.3% of the area exhibiting this pattern in 2018, while only ∼19.7% showed a unimodal pattern (Fig. 1a). A similar distribution was also observed in 2019 (Fig. S1). Two distinct photosynthetic peaks occurred in the first and second halves of the year, with average peak timings around March (DOY 81) and October (DOY 290), respectively (Fig. 1b).

**Fig. 1.**
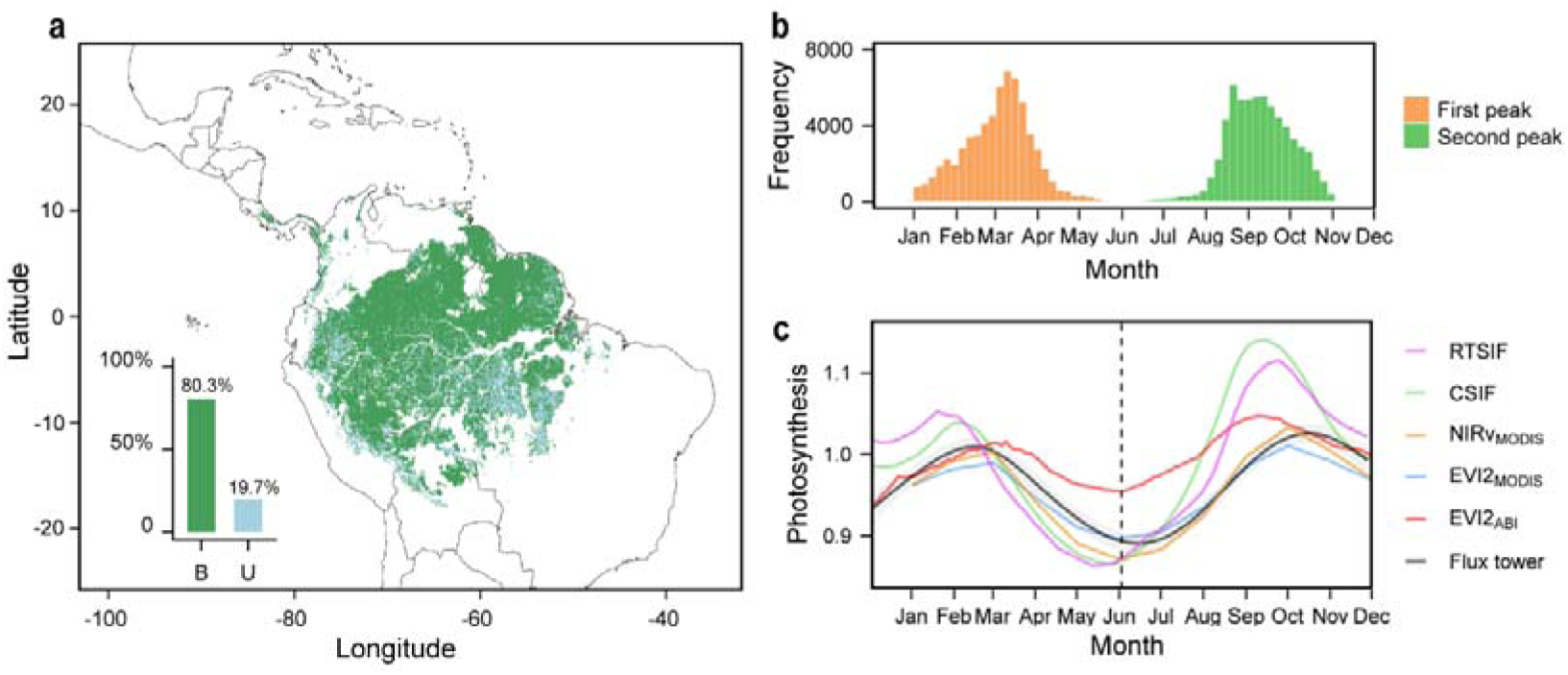
Bimodal photosynthetic pattern across the Amazon rainforest. **a**, Spatial distribution of photosynthetic seasonality in 2018. The blue series represents bimodal photosynthetic seasonality, whereas the green series represents unimodal photosynthetic seasonality. The bar chart illustrates the proportion of bimodal and unimodal photosynthetic seasonality across the region (with “B” indicating bimodal and “U” indicating unimodal). **b**, Distribution of the two photosynthetic peaks within the year. **c**, Intra-annual time series of different photosynthetic proxies. The vertical dashed line marks the half-year boundary (the 183rd day of the year).

To exclude potential biases from the detection algorithm, we further directly examined the bimodal pattern of photosynthesis using five remote sensing-based vegetation indices and flux tower-derived gross primary productivity (GPP). All datasets consistently revealed a pronounced bimodal seasonality in Amazon rainforest photosynthesis (Fig. 1c), which was evident across latitudinal gradients (Fig. S2) and further confirmed by flux tower observations at five out of six sites (Fig. S3). These findings show that bimodal photosynthetic seasonality is a pervasive feature in the Amazon rainforest, underscoring the need to disentangle climate sensitivities in each half of the year to better understand Amazon carbon dynamics under climate change.

### Differences in hydrothermal conditions and sensitivities

Over the period 2001-2020, the second half of the year is characterized by higher temperature and lower precipitation compared to the first half (Fig. 2a, b). Correspondingly, we found a significantly weaker S_T_ but stronger S_P_ in the second half of the year compared to the first half, based on four satellite-derived photosynthesis proxies and flux tower data (Fig. 2c, d). Temperatures consistently had a significant positive effect on photosynthesis in the first half of the year across all datasets, but this effect weakened and even turned negative in the second half based on flux tower observations (Fig. 2c). In contrast, precipitation showed no significant influence on photosynthesis in the first half of the year, yet exhibited a positive effect in the second half in satellite-derived datasets. Flux tower observations also revealed a positive effect of precipitation in the first half, which strengthened further in the second half (Fig. 2d).

**Fig. 2.**
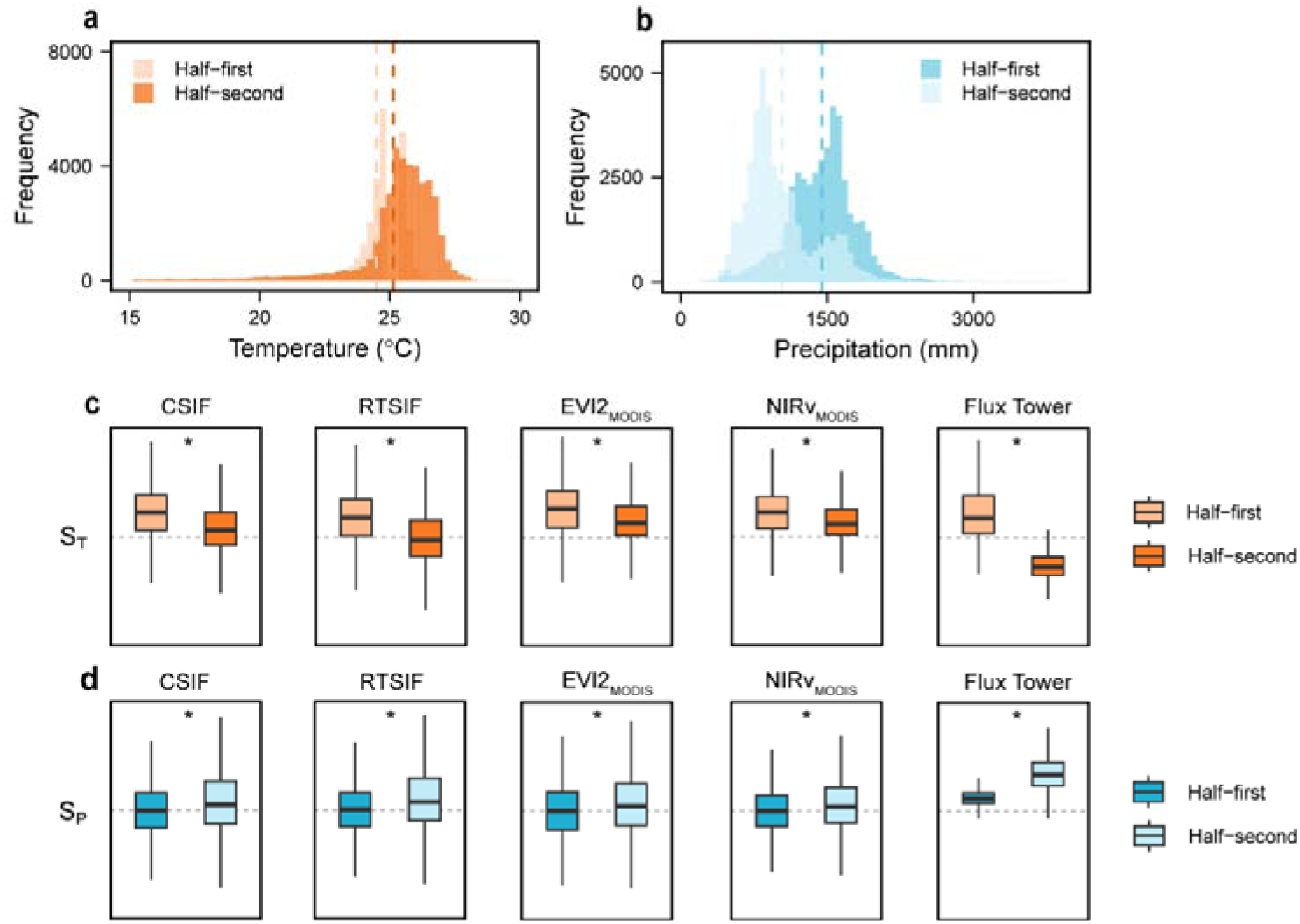
Differences in hydrothermal backgrounds and sensitivities between the two halves of the year. **a, b**, Differences in temperature (**a**) and precipitation (**b**) backgrounds. The vertical dashed lines mark the mean precipitation and temperature across the entire region for each half of the year. **c, d**, Comparisons of the overall S_T_ (**c**) and S_p_ (**d**) between the two halves of the year, with grey dashed lines marking zero. The boxes span from the first to the third quartile, with median values marked as the black lines in the boxes. The whiskers extend from the 5th to the 95th percentile. The asterisks indicate statistically significant differences (Student’s *t*-test; *p* < 0.05). To simplify, Half-first refers to the first half of the year, and Half-second refers to the second half of the year. These abbreviations are used consistently throughout the subsequent figure captions.

### Climatic drivers of hydrothermal sensitivities

To analyze the effects of temperature and precipitation on S_T_ and S_P_, we calculated the 20-year averages of temperature and precipitation for each half of the year and binned them into 10 × 10 percentile bins. Our findings revealed the hydrothermal drivers for S_T_ and S_P_, which remained consistent between both halves of the year (Fig. 3a, b). Specifically, S_T_ decreased with rising temperature and decreasing precipitation, while S_P_ exhibited the opposite pattern. We further examined the effects of air dryness (vapor pressure deficit, VPD) and soil moisture (SM) on photosynthesis and found that positive S_T_ and negative S_P_ were associated with lower VPD and higher SM (Fig. S4). These findings suggested that warmer and drier conditions frequently resulted in weaker S_T_ and stronger S_P_, whereas cooler and wetter conditions were linked with stronger S_T_ and weaker S_P_.

**Fig. 3.**
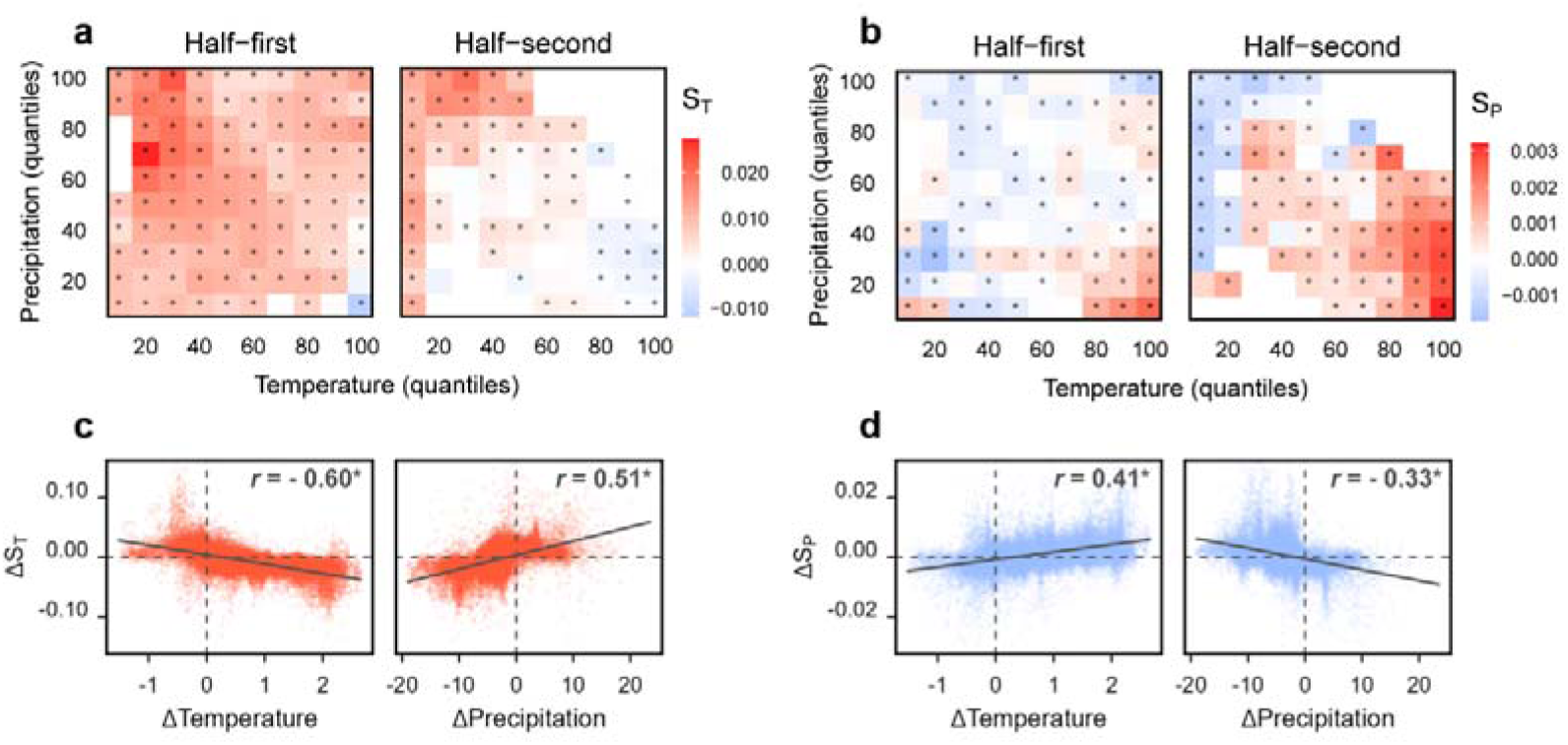
Hydrothermal mechanism driving variations in S_T_ and S_P_. **a, b**, S_T_ and S_P_ across hydrothermal bins in the two halves of the year. The asterisks represent the sensitivities which are significantly different from zero (*Wilcox*-test; *p* < 0.05) across all pixels for each bin. Bins containing fewer than 150 pixels were excluded. **c, d**, Relationship between changes in hydrothermal conditions and sensitivity. Delta (Δ) represents the change in value from the Half-first to the Half-second (Half-first minus Half-second). The solid lines indicate linear regression fits. The asterisks denote slopes which are statistically significant (Student’s *t*-test; *p* < 0.05). Temperature and precipitation units are °C and 100 mm, respectively.

To explain why hydrothermal sensitivities varied distinctly between the two halves of the year, we calculated the half-year differences for S_T_ and S_P_ (ΔS_T_, ΔS_P_), as well as for hydrothermal conditions (Δtemperature and Δprecipitation) for each pixel and detected their linkages. The results confirmed that increasing temperature (Δtemperature > 0) and decreasing precipitation (Δprecipitation < 0) significantly contributed to the reduced S_T_ (ΔS_T_ < 0) and rising S_P_ (ΔS_P_ > 0), and vice versa (Fig. 3c, d).

### Temporal change in hydrothermal sensitivities

To examine and compare temporal changes in the photosynthetic response to hydrothermal variations between the two halves of the year, we applied a moving window method to detect shifts in hydrothermal sensitivities (See Methods). We observed a faster warming rate alongside greater changes in the proportions of positive S_T_ and S_P_ in the second half of the year compared to the first half. During 2001-2020, significant and asymmetric warming trends were observed between the two halves of the year, with rates of +0.18D·decade^−1^ in the first half and +0.32D·decade^−1^ in the second half (Fig. 4a). Conversely, insignificant trends were observed for regional average annual precipitation across the Amazon rainforest (Fig. 4b). For hydrothermal sensitivities during 2001-2020, the proportion of positive S_T_ showed a declining trend, with a sharper decrease in the second half (from 72.4% to 25.1%, −5.3%·year^−1^) than in the first half (from 87.9% to 52.6%, −3.0%·year^−1^) (Fig. 4c). On the other hand, the proportion of positive S_p_ exhibited an increasing trend, with a sharper elevation also in the second half (from 44.4% to 68.5%, +2.6%·yearD¹) compared to the first half (from 46.9% to 51.4%, +0.6%·year^−1^) (Fig. 4d). Moreover, significant correlations were observed between temperature and the proportions of positive hydrothermal sensitivities in both halves of the year (first half: *r* = −0.77 for S_T_, *r* = 0.74 for S_P_, both *p* < 0.05; second half: *r* = −0.97 for S_T_, *r* = 0.96 for S_P_, both *p* < 0.05). In contrast, precipitation changes showed no significant contribution (*p* > 0.05). It should be noted that we only present results using a 10-year moving window, as varying the window width had minimal impacts on the trends. The results from ridge regression also exhibited similar outcomes (Fig. S5).

**Fig. 4.**
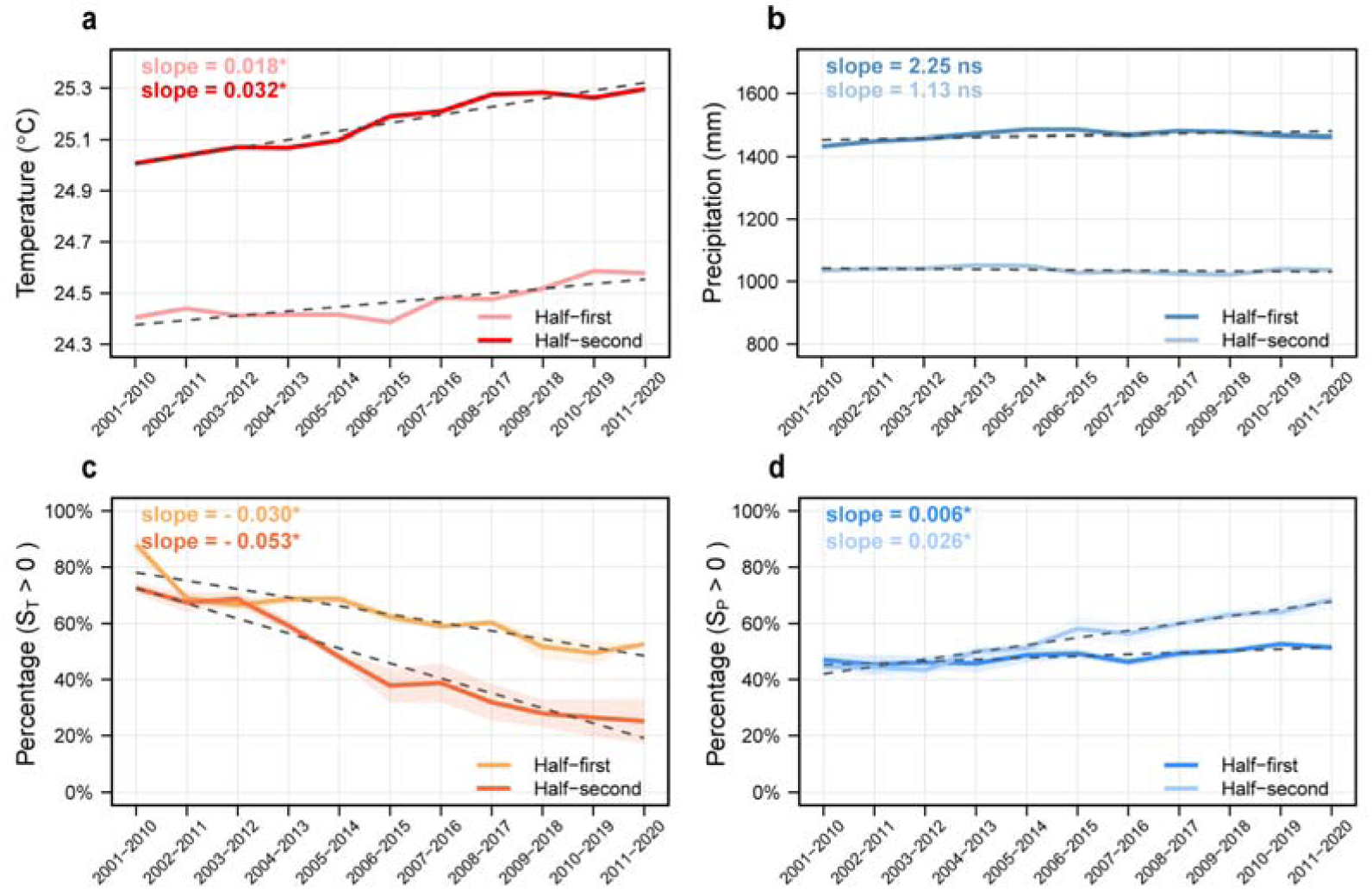
Temporal changes in hydrothermal conditions and sensitivities. **a, b**, Temporal changes in the regional average temperature (**a**) and precipitation (**b**). **c, d**, Proportions of positive S_T_ (**c**) and S_P_ (**d**) over time. The solid lines in **c** and **d** indicate the multi-dataset average, with the shaded area representing one standard deviation on either side of the mean. The dashed lines in a-d represent the linear regression fits, and the asterisks represent the slopes that are statistically significant (Student’s *t*-test; *p* < 0.05).

### T_opt_ and Year_tipping_ in the Amazon rainforest

The significant decrease in the proportion of positive S_T_ mentioned above suggests that a substantial portion of the Amazon rainforest has surpassed its interannual T_opt_ (where S_T_ shifts from positive to negative) during the past two decades. We found that the Amazon rainforest experienced a widespread decrease in S_T_ over the past two decades, with reductions of 73.7% in the first half of the year and 84.4% in the second half, responding to an overall warming trend (88.6% in the first half of the year and 98.9% in the second half). Subsequently, we used quadratic functions to identify the T_opt_ in the pixels where temperature increased and S_T_ decreased over the past two decades (See Methods).

Our findings revealed a 0.49□ higher T_opt_ on average in the second half of the year (25.63 ± 0.02□) than in the first half (25.14 ± 0.02□) (*p* < 0.05, Fig. 5a). Temperature was proven to be the primary factor influencing T_opt_ rather than other climatic factors or baseline photosynthetic capacity, as determined through random forest analysis (Fig. S6). Regression analysis further demonstrated that T_opt_ increased significantly with rising temperature in both halves of the year (Fig. 5b). By combining historical (ERA-Land) and future (ESMs) temperatures, we investigated the tipping year (Year_tipping_) when T_opt_ emerges (temperature effect reversed) for each half of the year (Fig. 5c, d). We found that a substantial portion of the area experienced a reversal of climate warming benefits over the past 20 years, with the more intense reversal occurring in the second half of the year. By 2017 and 2014, half of the areas had already reached T_opt_ for the first and second halves of the year, respectively. We further simulated Year_tipping_ from 2020 to 2100 and compared the intensity of reversals across three Shared Socioeconomic Pathways (SSP) scenarios (see Methods). The overall intensity of the reversal is expected to increase with the rate of climate warming. Under the most critical scenario (SSP585), more than 90% of the areas are expected to reach T_opt_ in the first and second halves of the year by 2035 and 2031, respectively.

**Fig. 5.**
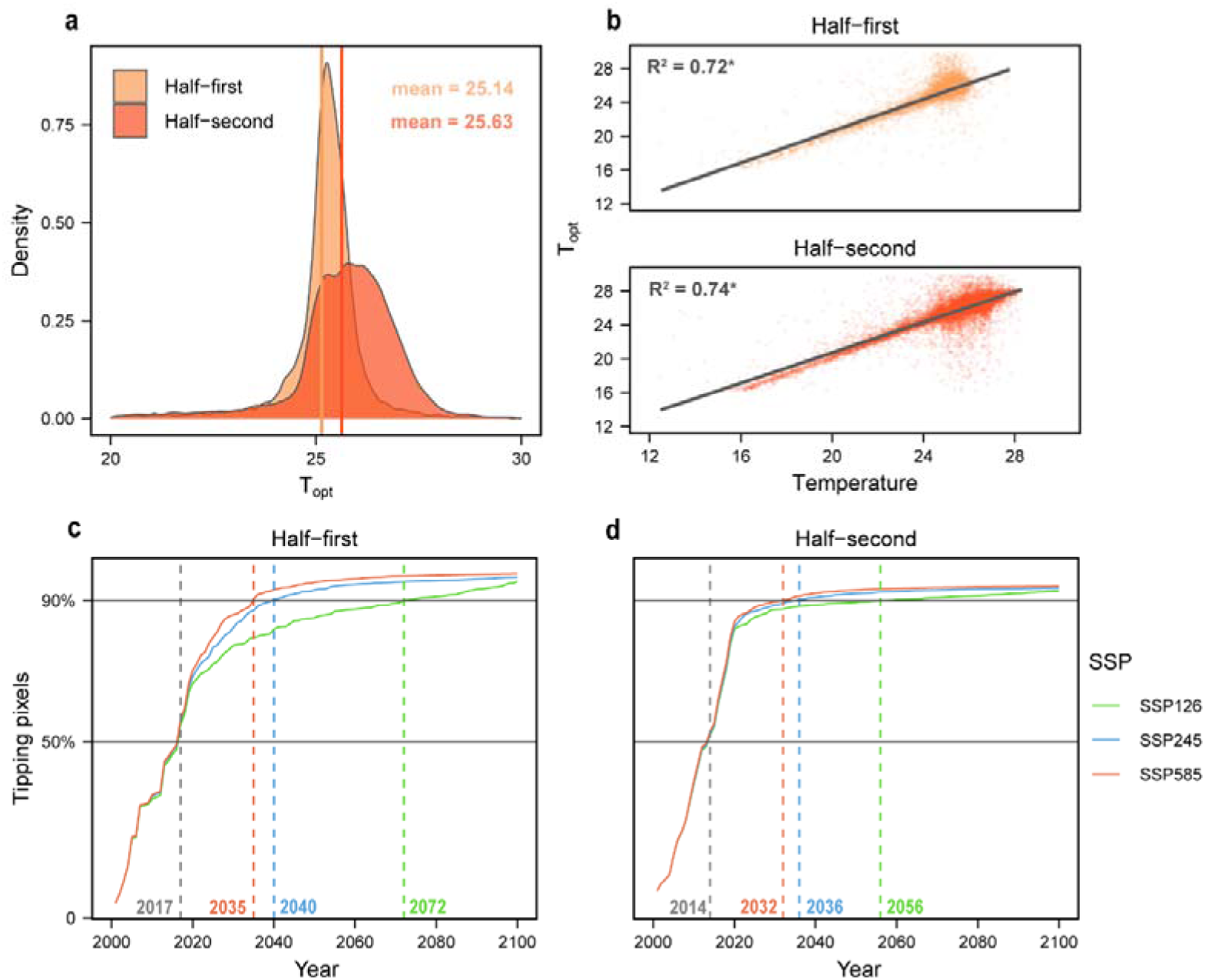
Reversal pattern of temperature effects. **a**, Difference of T_opt_ between the two halves of the year. The vertical solid lines indicate the mean T_opt_ for each half of the year. **b**, Linear relationship between T_opt_ and the temperature background. The asterisks represent the slopes that are statistically significant (Student’s *t*-test; *p* < 0.05). **c, d**, Trajectory of the tipping pixels under different Shared Socio-economic Pathways (SSPs). The horizontal solid lines mark the 50% and 90% thresholds, and the vertical dashed lines denote the years when these thresholds are reached under different SSPs.

### Reversed temperature effects on photosynthesis

To assess differences in the temporal trends of photosynthetic capability between the first and second halves of the year, we mapped the interannual photosynthetic trends for each half of the year across two periods (2001–2010 and 2011–2020). We found that the photosynthetic trend reversal (from positive slope to negative slope) mainly occurred in the northwestern Amazon in the first half of the year, while it was observed in the central and northern Amazon in the second half (Fig. 6a, b), coinciding with areas where the temperature effect reversed during 2001-2020 (the blue areas in Fig. S7). Compared to the first half of the year, the second half exhibits a more pronounced decline in photosynthesis, leading to changes in the intra-annual photosynthesis proportion. With climate warming, the proportion of photosynthesis in the second half of the year decreased in approximately 76.6% of areas during 2001-2020, contributing significantly to the annual photosynthetic decline (Fig. 6c). These findings demonstrate that climate warming has severely harmed photosynthetic activity and greatly degraded the overall photosynthetic capability of the Amazon rainforest.

**Fig. 6.**
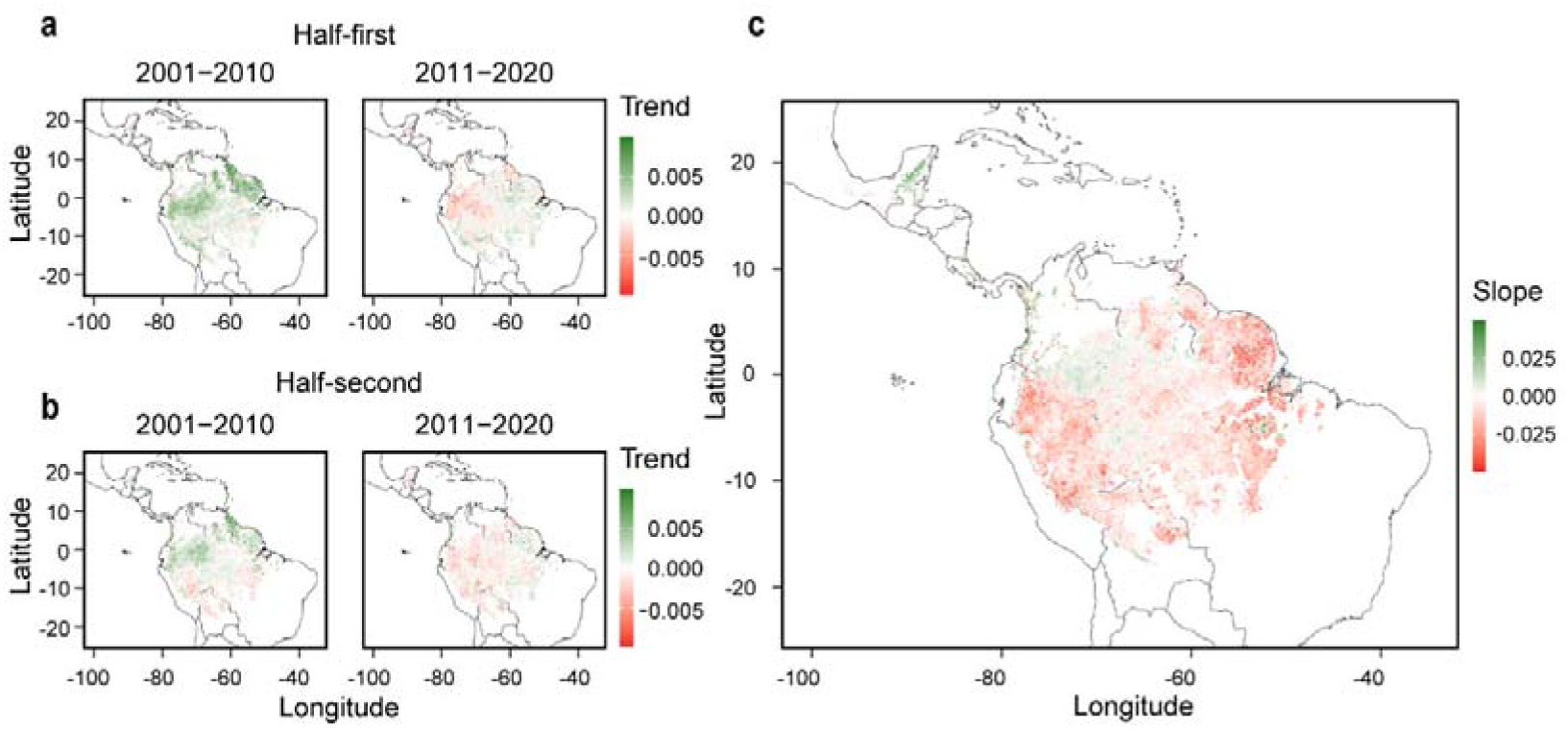
Spatial pattern of reversed temperature effects on photosynthesis. **a, b**, Photosynthesis trends in two periods for each half of the year, calculated by regressing CSIF against year. **c**, Change in the proportion of Half-second photosynthesis, with the slope derived from the regressing proportion of Half-second photosynthesis against annual temperature. The green series indicates positive values, whereas the red series indicates negative values. The bar chart illustrates the proportion of positive and negative values (“P” represents positive, “N” represents negative).

## Discussion

### Bimodal photosynthetic pattern in the Amazon rainforest

Vegetation physiological dynamics are tightly coupled with seasonal climate variability. In temperate and boreal forests, annual energy limitations—primarily temperature and radiation—produce a unimodal pattern of photosynthetic activity. In contrast, bimodal photosynthetic seasonality is frequently observed in dry land and tropical rainforests, reflecting vegetation’s adaptive responses to asynchronous fluctuations in energy and water availability^14,23,42–47^. For example, in Mediterranean ecosystems, plant growth is initially stimulated by rising spring temperatures and replenished soil moisture. However, during late spring and summer, extreme heat and severe drought suppress photosynthetic activity until reactivated by late-summer or autumn rainfall, thus generating bimodal growing season^42,43^. Similarly, in tropical rainforests, the biannual migration of the solar zenith angle results in two periods of reduced solar radiation, temporarily suppressing photosynthetic activity^14,16,20,21,23^. This radiative pattern underpins the emergence of two distinct photosynthetic peaks across the year. In the Amazon rainforest, this bimodal seasonality is further shaped by contrasting hydrothermal conditions between the first and second halves of the year, which likely serve as a primary driver of different intra-annual photosynthetic responses.

### Intra-annual difference in hydrothermal sensitivities

Based on the intra-annual bimodal photosynthetic pattern divided by half-year intervals, we found that temperature sensitivity (S_T_) was generally weaker, whereas precipitation sensitivity (S_P_) was stronger in the second half of the year compared to the first half in the Amazon rainforest. We attributed these intra-annual differences in S_T_ and S_P_ to hydrothermal conditions. In the warmer and drier second half of the year, elevated temperatures and reduced precipitation impose greater constraints on photosynthesis. Higher temperatures exacerbate water stress by increasing evapotranspiration, prompting trees to close their stomata to minimize water loss^48–50^. This stomatal closure directly reduces COD intake, thereby decreasing photosynthetic rates and weakening the temperature-driven enhancement of photosynthesis, as reflected in the lower S_T_. Additionally, lower precipitation increases the risk of gas embolism in water transport systems, impairing the movement of water from roots to leaves^51,52^. These disruptions limit canopy photosynthesis and hinder nutrient uptake and translocation within the tree, further suppressing photosynthetic activity^53^. Consequently, tree growth in the second half of the year is more susceptible to water limitation, as indicated by the higher S_P_ compared to the first half.

In contrast, in the first half of the year, characterized by lower temperatures and higher precipitation, moderate warming can enhance enzymatic activity and metabolic processes associated with photosynthesis^54^, leading to an increase in S_T_. Ample precipitation ensures sufficient water availability for stomatal opening and nutrient absorption, reducing water limitations on photosynthesis and resulting in a lower S_P_.

In summary, the warmer and drier conditions in the second half of the year amplify water stress, reducing S_T_ while increasing S_P_. Conversely, the cooler and wetter conditions in the first half enhance temperature-driven photosynthetic gains while mitigating precipitation-related constraints. These findings underscore the crucial role of hydrothermal conditions in co-regulating intra-annual photosynthetic responses by influencing COD intake, water transport, and nutrient translocation within trees in the Amazon rainforest.

### Asymmetric temporal change in hydrothermal sensitivities

Building upon the observed intra-annual differences in S_T_ and S_P_, our analysis further revealed asymmetric temporal trends in these sensitivities over the past two decades. Specifically, while S_T_ remained positive in both halves of the year around 2001, the regions with positive S_T_ where rising temperature effectively enhanced photosynthesis have been steadily shrinking since then. This shrinkage was more pronounced in the second half of the year compared to the first half. In contrast, S_P_ showed an increasing trend over this period, suggesting that the regions where photosynthesis is limited by precipitation have expanded, particularly in the second half of the year. These findings indicate a transition from temperature limitation to precipitation limitation, especially during the second half of the year in the Amazon rainforest.

To explain these asymmetric changes in S_T_ and S_P_ over time, we examined trends in temperature and precipitation from 2001 to 2020. Our results indicate that precipitation levels have remained relatively stable in both halves of the year. However, the second half of the year has experienced a more rapid warming trend, despite temperature increases in both halves of the year. Therefore, the faster warming rate in the second half of the year may have exacerbated evapotranspiration, intensifying soil moisture depletion. This reduced photosynthetic rates by amplifying stomatal closure and triggering gas embolism in the water transport systems, disrupting the movement of water and nutrients^52,53^. These factors likely contributed to the more rapid decline in areas with positive S_T_ and the more pronounced increase in areas with positive S_P_ compared to the first half of the year.

Overall, our findings provide robust evidence that the Amazon rainforest is transitioning from temperature limitation to precipitation limitation, particularly in the second half of the year. This underscores a growing dependence of vegetation physiological processes on water availability. However, this increasing demand appears challenging to meet due to relatively stable precipitation levels. Additionally, intensifying precipitation seasonality and a prolonged dry season are likely to exacerbate water stress during critical growth periods, collectively posing heightened threats to plant growth and increasing drought-induced tree mortality^37,39,55–57^. Water stress-induced growth limitations could further disrupt the regional water cycle through altered transpiration, amplifying climate-related risks^37,58,59^. Therefore, this pivotal transition has profound implications for sustaining forest productivity and stability under future climate change. As such, continuous and long-term monitoring is crucial for understanding and predicting how hydrothermal variations may reshape carbon-water cycling in the Amazon rainforest.

### Intra-annual difference in thermal acclimation

Photosynthetic activity in tropical forests, including the Amazon, exhibits nonlinear responses to temperature, typically following a convex curve with an optimal temperature threshold (T_opt_). Understanding T_opt_ is crucial for evaluating how these ecosystems will respond to changing hydrothermal conditions, particularly as global temperatures continue to rise.

Recent studies suggest that the Amazon rainforest may already be operating near or beyond its T_opt_^27,28^, raising concerns about the forest’s ability to maintain photosynthetic efficiency under climate warming. Given our findings that vegetation hydrothermal sensitivities varied distinctly between the two halves of the year, it is likely that thermal acclimation of vegetation also differs accordingly. Therefore, to gain a deeper understanding of the relationship between vegetation photosynthesis and climate, we separately investigated T_opt_ for each half of the year. We found that the second half of the year exhibited a significantly higher T_opt_ than the first half.

This result aligns with the beneficial acclimation hypothesis, which suggests that organisms can adjust their physiological processes to optimize performance in response to thermal conditions^60,61^. Given that temperatures in the second half of the year are generally higher than in the first half, it is expected that the optimal temperature for photosynthesis would also increase^62^. The higher T_opt_ observed in the second half of the year implies that the Amazon rainforest can acclimate to increased seasonal temperatures by shifting its optimal photosynthetic temperature range and thus assimilate more carbon during this period. However, although the second half of the year has a higher T_opt_, it reaches this optimal temperature earlier than the first half. This scenario could be attributed to the more rapid warming in the second half of the year, which may amplify hydrothermal stress (as mentioned earlier) and hinder further thermal acclimation, thereby accelerating the attainment of T_opt_.

### Asymmetric temporal change in photosynthesis

Under warming, the temperature effect on photosynthesis is projected to reverse across Northern Hemisphere ecosystems^63^. Herein, we found that the Amazon rainforest is currently undergoing an intensified reversal of the temperature effect, which has led to a widespread decline in photosynthesis. This decline impairs the Amazon rainforest’s capability to assimilate COD, adding to the growing evidence of its weakening role as a global carbon sink. Since photosynthesis is key to energy flow and matter cycling, its decline may also challenge the Amazon rainforest in maintaining its ecological structure and function^64,65^.

To clarify the consequences of the reversed temperature effects coupled with the asymmetric temporal decline in S_T_, we directly analyzed the photosynthesis trend in the Amazon rainforest during the periods 2001-2010 and 2011-2020. We revealed a widespread reversal in photosynthesis during these two halves of the year, coinciding with regions where the temperature effects reversed. A more pronounced decline in photosynthesis was observed in the second half of the year, resulting in a reduced proportion of annual photosynthesis occurring in this period. This shift in the seasonal contribution of carbon uptake by the two halves of the year may exacerbate the intra-annual imbalance in carbon dynamics within the Amazon rainforest.

Many growth processes, such as leaf-out, flowering, and fruit production, are closely tied to the seasonal dynamics of carbon acquisition and allocation^66–68^. This shift in seasonal carbon uptake could perturb the temporal coordination or efficacy of growth rhythm, misaligning plant phenology with the life cycles of dependent organisms. Furthermore, this may result in potential intra-annual redistribution of biological resources, triggering significant shifts in community structure within the Amazon rainforest. Such changes could destabilize the entire ecosystem when coupled with thermal stress and other ongoing threats, such as climatic disturbances, deforestation, fires, and forest fragmentation, all of which exacerbate reductions in photosynthetic productivity^9,11,12,69^. Therefore, targeted strategies are crucial for mitigating the adverse effects of climate change and sustaining forest productivity, particularly in the second half of the year, to address both seasonal and long-term ecological stability of the Amazon rainforest.

## Conclusion

By integrating multiple datasets with direct observations and phenological extraction methods, our study not only provides strong evidence that the Amazon rainforest exhibits widespread bimodal photosynthetic seasonality across the first and second halves of the year but also reveals asymmetric photosynthetic responses to hydrothermal variability between these two periods. We observed a weaker temperature limitation but a stronger precipitation limitation on photosynthesis in the second half of the year compared to the first half in the Amazon rainforest. Under climate warming, photosynthesis in the Amazon rainforest is shifting from temperature limitation to precipitation limitation, particularly during the second half of the year. Although optimal temperatures for photosynthesis are higher in the second half, they are reached earlier, leading to a sharper decline in productivity and a reduced contribution to total annual photosynthesis. Our findings not only provide further evidence of the increasing dominance of water limitation but also highlight pronounced seasonal asymmetry in photosynthetic responses to climatic stressors in the Amazon rainforest. These results enhance our understanding of tropical forest growth under climate change and offer important insights for improving vegetation models and increasing the accuracy of future ecosystem projections.

## Methods

### Study area

Our study focused on the Amazon region, where evergreen broadleaf forests are predominantly distributed. The study area was defined using the MODIS Collection 6.1 Land Cover Product (MCD12C1), at a spatial resolution of 0.05°, and included pixels classified as evergreen broadleaf forests within the tropical Americas (23.5° S to 23.5° N, 40° W to 100° W). Although a small portion of these forests is located in Central America, the vast majority lies within the Amazon basin. Therefore, for simplicity, we collectively refer to this region as the Amazon rainforest in this study.

### RTSIF and CSIF

As a direct measure of photons intercepted by chlorophyll, solar-induced chlorophyll fluorescence (SIF) serves as a robust proxy for vegetation photosynthesis and is widely used in studying the Amazon rainforest^30,70–72^. In this study, two sets of spatiotemporally continuous clear-day SIF data from 2001–2020 were used: the reconstructed TROPOspheric Monitoring Instrument (TROPOMI) SIF (RTSIF)^73^ and the Contiguous Solar-Induced Fluorescence (CSIF)^74^. The RTSIF was reconstructed based on TROPOMI SIF measurements using machine learning with a spatial resolution of 0.05° at each 8-day interval. The CSIF dataset was generated at a spatial resolution of 0.05° with a 4-day time step by training a neural network with SIF from the Orbiting Carbon Observatory-2 (OCO-2). Importantly, the CSIF dataset avoids using hydrothermal data, thereby mitigating artificially induced hydrothermal-vegetation growth coupling effects. Therefore, the results presented are primarily based on CSIF to avoid excessive redundancy.

### NIRv and EVI2

The Near-Infrared Reflectance of Vegetation (NIRv) and the Two-band Enhanced Vegetation Index (EVI2), both derived from red and near-infrared reflectance, are reliable proxies for photosynthesis due to their strong resistance to saturation and effectiveness in capturing canopy structure^75–77^. In this study, we derived long-term NIRv and EVI2 from the MODIS Collection 6.1 MCD43C4 Nadir BRDF-Adjusted Reflectance (NBAR) product, which provides a spatial resolution of approximately 0.05° and a daily temporal resolution, covering the period from 2001 to 2020. This product leverages a semi-physical BRDF model to minimize the influence of solar-view geometry variations on vegetation reflectance, which often leads to inconsistent conclusions^33^. The MCD43C4 data quality has been evaluated across globally representative locations and time periods through comparisons with reference *in-situ* and other independent datasets^33,78,79^. In our study, we utilized observations classified as mixed to best quality and subsequently computed the monthly medians of the derived indices.

Additionally, we derived EVI2 from the Advanced Baseline Imager (ABI) onboard the Geostationary Operational Environmental Satellite 16 (GOES-16), positioned over the equator at approximately 70° W. The ABI scans the entire Amazon region with a sub-hourly revisit frequency: every 15 minutes in 2018 and every 10 minutes in 2019, at a spatial resolution of approximately 1.0 km^80^. Compared to polar-orbiting satellites such as MODIS or VIIRS, ABI has been shown to provide 21-35 times more clear-sky observations per month and detects more than three times the seasonal changes in the Amazon rainforest^81^.

### ABI NBAR EVI2

To generate a high-quality ABI NBAR EVI2 time series, we first extracted red and near-infrared band reflectance from the ABI Multi-Angle Implementation of Atmospheric Correction (MAIAC) surface reflectance product, along with associated quality assurance (QA) metrics and solar and viewing geometry angles, for the period from January 1, 2018, to December 30, 2019. Next, all available clear-sky ABI reflectance within non-overlapping 3-day windows was used to construct the Ross-Thick-Li-Sparse-Reciprocal (RTLSR) model, which has long been employed for generating NBAR products for both VIIRS and MODIS^82^. The selection of a 3-day window was aimed at balancing the availability of ABI observations with the stability of surface conditions^83,84^ Only high-quality BRDF retrievals that met specific criteria during all 3-day rolling periods were retained^84^. Finally, the resultant high-quality NBAR data at the red and near-infrared bands were used to calculate EVI2, representing the photosynthetic status for each 3-day period in 2018 and 2019.

### Flux tower data

Ground-based flux observations were used to complement analyses of photosynthetic seasonality and hydrothermal sensitivities in the Amazon rainforest. Hourly or daily data of gross primary productivity (GPP), precipitation, and temperature were acquired from the FLUXNET2015, LBA-ECO CD-32 Flux Tower Network, AMERIFLUX, and Amazon Tall Tower Observatory (ATTO) project^85^. We first selected sites within the study region classified as evergreen broadleaf forest. We then selected years in which the number of days with valid data exceeded 80% in both halves of the year for each site. Notably, for sites that appeared in multiple datasets, we selected the dataset with the most available years. Details of the flux sites are provided in Table S1.

### Climate data

Monthly data for radiation, soil moisture, and dew point temperature spanning the period 2001–2020, as well as temperature data from 1950 to 2020, were obtained from the ERA5-Land reanalysis dataset at a spatial resolution of 0.1°. The vapor pressure deficit (VPD) was calculated using the Magnus formula, incorporating temperature and dew point temperature^86^. Precipitation data were sourced from the Climate Hazards Group InfraRed Precipitation with Station data (CHIRPS), which offers monthly precipitation data at a spatial resolution of 0.05°.

### Earth System Model (ESM) data

Monthly temperature data from eight Coupled Model Intercomparison Project Phase 6 (CMIP6) models were utilized, spanning the period from 2001 to 2100. This dataset includes historical data (2001–2014) and simulated projections (2015–2100) under three Shared Socioeconomic Pathways (SSP126, SSP245, and SSP585). The eight models used are: CanESM5, ACCESS-ESM1-5, BCC-CSM2-MR, CMCC-CM2-SR5, INM-CM4-8, IPSL-CM6A-LR, MPI-ESM1-2-LR, and NorESM2-LM.

All gridded time series datasets were aggregated to a resolution of 0.1° to ensure consistency of spatial resolution and then computed to create two half-year averages for each year and each pixel, respectively. Note that the extraction of photosynthetic peak timings is based on the aggregated ABI EVI2 time series.

### Analysis

#### Examination of bimodal photosynthetic pattern

We adapted a spline-based land surface phenology (LSP) detection algorithm from the operational MODIS LSP product^74^ to extract the timing of peak photosynthesis from the ABI NBAR EVI2 time series in the Amazon rainforest during 2018 and 2019. First, we cleaned the high-quality ABI NBAR EVI2 time series by removing anomalously high or low values using a Z-score-based outlier detection method (±2σ). The data were then smoothed and gap-filled using a combination of moving averages, moving medians, and Savitzky-Golay filters with a window size of 7 (21 days), which are the standard procedure during LSP detection process. The resulting time series was further fitted using a quality assurance (QA)-weighted penalized cubic smoothing spline. Candidate peaks in the spline-fitted time series were identified as local maxima, defined by zero-crossings in the first derivative. To ensure data reliability, we applied a series of ecological filters to exclude implausible peaks^87^, including: (1) a minimum EVI2 amplitude of 0.1 for annual seasonal variation; (2) a peak EVI2 exceeding 0.5 in evergreen broadleaf forest pixels; (3) a minimum 60-day interval between adjacent peaks; (4) a difference between peak and minimum EVI2 exceeding 35% of the annual EVI2 amplitude; and (5) a peak-to-peak EVI2 difference below 0.1. The final output was the timing of peak photosynthesis, expressed as the day of year (DOY). To reduce potential biases from the phenology detection algorithm, we also examined seasonal variations in photosynthesis using five satellite-derived photosynthetic proxies. We calculated the multi-year average of each photosynthetic proxy at each time interval within the year across the entire study area and along latitudinal gradients, using 2° intervals.

#### Calculation of S_T_ and S_P_

To establish interpretable and comparable indicators, linear regression was performed using the half-year mean photosynthesis proxies, as well as temperature and precipitation for each pixel. The slope obtained from each regression was defined as the sensitivity to temperature (S_T_) or precipitation (S_P_), indicating the change in photosynthesis intensity per 1 □ or 100 mm increase in temperature or precipitation, respectively. Positive sensitivities indicate an enhancement of photosynthesis by the increase in temperature or precipitation, and vice versa. Furthermore, we employed ridge regression to exclude the effects of VPD and radiation which have been shown to strongly influence photosynthesis in the Amazon rainforest, and account for bias in coefficient estimates caused by potential multicollinearity among variables. By introducing a penalty term (lambda, λ) into the minimized residual equation, ridge regression can reduce the variance of the coefficient estimate, making it more stable and less sensitive to other correlated predictors, thus improving the overall robustness of the model against multicollinearity. Note that the coefficient estimates from ridge regression are biased and generally smaller than those from linear regression due to the different selection of penalty terms in each ridge regression^88^. Therefore, the coefficient estimates from different ridge regressions are not directly comparable, while their signs (+/−) remain meaningful. Additionally, we also explored the effects of VPD and SM on photosynthesis as an additional analysis for water conditions, in addition to precipitation. Given the limited observation years per flux site, data from all sites were pooled, and bootstrap resampling (1,000 iterations) was performed to estimate hydrothermal sensitivities for each half of the year.

#### Temporal trend detection

A 10-year moving window was employed to explore temporal trends of hydrothermal conditions and sensitivities from 2001 to 2020. Both linear regression and ridge regression were performed to obtain S_T_ and S_P_ at each pixel in each moving window. For each moving window, we calculated the areal percentages of the positive sensitivities (S_T_ > 0 and S_p_ > 0), allowing for integration across different datasets and providing valuable insights into areal changes. Furthermore, we calculated the averages of temperature and precipitation in each moving window and detected their linkages with the percentages of positive sensitivities. We also employed different widths (8- and 12-year) of the moving windows to strengthen our temporal trend results.

#### Derivation of T_opt_ and Year_tipping_

We initially calculated S_T_ (for each pixel) in two periods (2001-2010 and 2011-2020) and classified all pixels into four types (PP, PN, NN, NP) based on changes in the positive-negative relationship. For example, a pixel with a positive S_T_ in 2001-2010 and a negative S_T_ in 2011-2020 was labeled as “PN”. We detected the temporal linear trend of S_T_ and temperature for each pixel and selected the pixels with increasing temperature and decreasing S_T_. We combined them with the four types above and excluded a few contradictory conditions (decreasing S_T_ with “NP”). Finally, we identified three types of temporal variations in S_T_ (decrease-NN, decrease-PN, decrease-PP). Notably, only in the “decrease-PN” type can we confirm a reversal of the temperature effect on photosynthesis during 2001-2020, while reversals may occur during or outside this timeframe in the other two situations. We subsequently fitted a quadratic function (Equation 1) following previous studies^62,63,89,90^ for each pixel within the three types above:

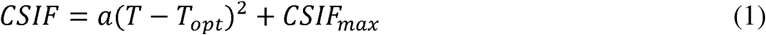

where *a* represents the direction and extent of the parabola. We identified the cases with *a*<0 as valid fitting due to the natural down-concave shape of the response curve of photosynthesis to temperature. T_opt_ is the temperature at which CSIF_max_ emerges, and we excluded outliers below the 1st percentile and above the 99th percentile. To ascertain the robustness of the T_opt_ retrievals, we further derived T_opt_ using two alternative methods. First, we identified the tipping period at the 10-year moving window when S_T_ shifts from positive to negative for the first time (Method2)^63^. Within the tipping period, the mean temperature was defined as T_opt_. Second, the T_opt_ was defined as the temperature where CSIF was maximized during 2001-2020 within the “decrease-PN” type (Method3). Finally, the results from the two alternative methods yielded similar outcomes (Fig. S8).

We integrated historical temperatures from ERA5-Land (1950-2020) and ESM temperatures averaged from 8 CMIP6 models (2021-2100) under three Shared Socioeconomic Pathways (SSP126, SSP245, and SSP585). We calibrated the temperatures from ESM models based on the baseline of 2001-2020 for the ERA5-Land data and ESMs in three Shared Socio-economic Pathways (SSP126, SSP245, SSP585) following a reference^91^. Specifically, we calculated the half-year average temperatures for ERA5-Land data 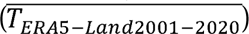 and ESMs outputs 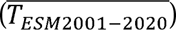 during 2001-2020 at each pixel, respectively. Leveraging the bias between the two temperature datasets, the calibrated temperature was generated for each pixel and each half of the year as follows:

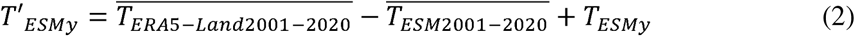

where *y* represents the specific year, *T_ESM_* is the original averaged temperature from 8 ESMs, and *T’_ESM_* is the calibrated averaged temperature from ESMs. It is important to note that biases arise from modeling mechanisms and are not consistent system errors. Thus, perfectly aligning future climate data from ESMs with historical ERA5-Land data is impossible. To minimize uncertainty in Year_tipping_ derivation, we calculated the absolute difference between temperature and T_opt_ for each half of the year from 1950 to 2100. Year_tipping_ was identified as the year with the minimum absolute difference from 1950 to 2020, 2001 to 2020, and 2001 to 2100 for the decrease-NN, decrease-PN, and decrease-PP types, respectively, and we excluded cases where the minimum absolute difference was greater than 0.2 °C.

## Data availability

The MCD43C4 product was obtained from the NASA Earthdata platform (https://e4ftl01.cr.usgs.gov), and the MCD12C1 dataset is available from the LP DAAC Data Pool (https://lpdaac.usgs.gov/tools/data-pool). CSIF and RTSIF datasets can be accessed via the National Tibetan Plateau Data Center (https://data.tpdc.ac.cn/zh-hans/data). ABI data are available from the NASA GeoNEX portal (https://data.nas.nasa.gov/geonex/data.php). Climate variables, including temperature, radiation, soil moisture, and dew point temperature, were retrieved from the ERA5 reanalysis dataset (https://cds.climate.copernicus.eu/datasets). Precipitation data were obtained from the CHIRPS dataset (https://chc.ucsb.edu/data). The FLUXNET2015 dataset is available at https://fluxnet.org/data/fluxnet2015-dataset. The LBA-ECO CD-32 flux tower network data can be accessed through the ORNL DAAC (https://daac.ornl.gov/LBA/guides/CD32_Fluxes_Brazil.html). The AMERIFLUX dataset is available at https://ameriflux.lbl.gov/data/download-data. The ATTO data is requested at https://www.attodata.org/ddm/search. Earth system model (ESM) outputs from CMIP6 were downloaded via the ESGF data portal (https://esgf-node.ipsl.upmc.fr/search/cmip6-ipsl).

## Code availability

Codes to analyze the data are available from https://doi.org/10.6084/m9.figshare.29046362.v1.

---

**Fig. S1.**
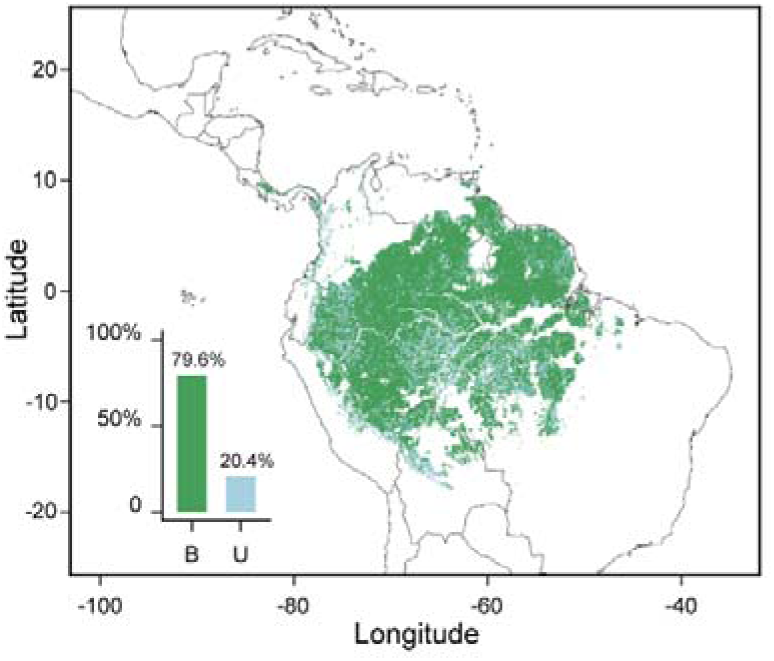
Spatial distribution of photosynthetic seasonality in 2019. The blue series represents bimodal photosynthetic seasonality, whereas the green series represents unimodal photosynthetic seasonality. The bar chart illustrates the proportion of bimodal and unimodal photosynthetic seasonality across the region (with “B” indicating bimodal and “U” indicating unimodal).

**Fig. S2.**
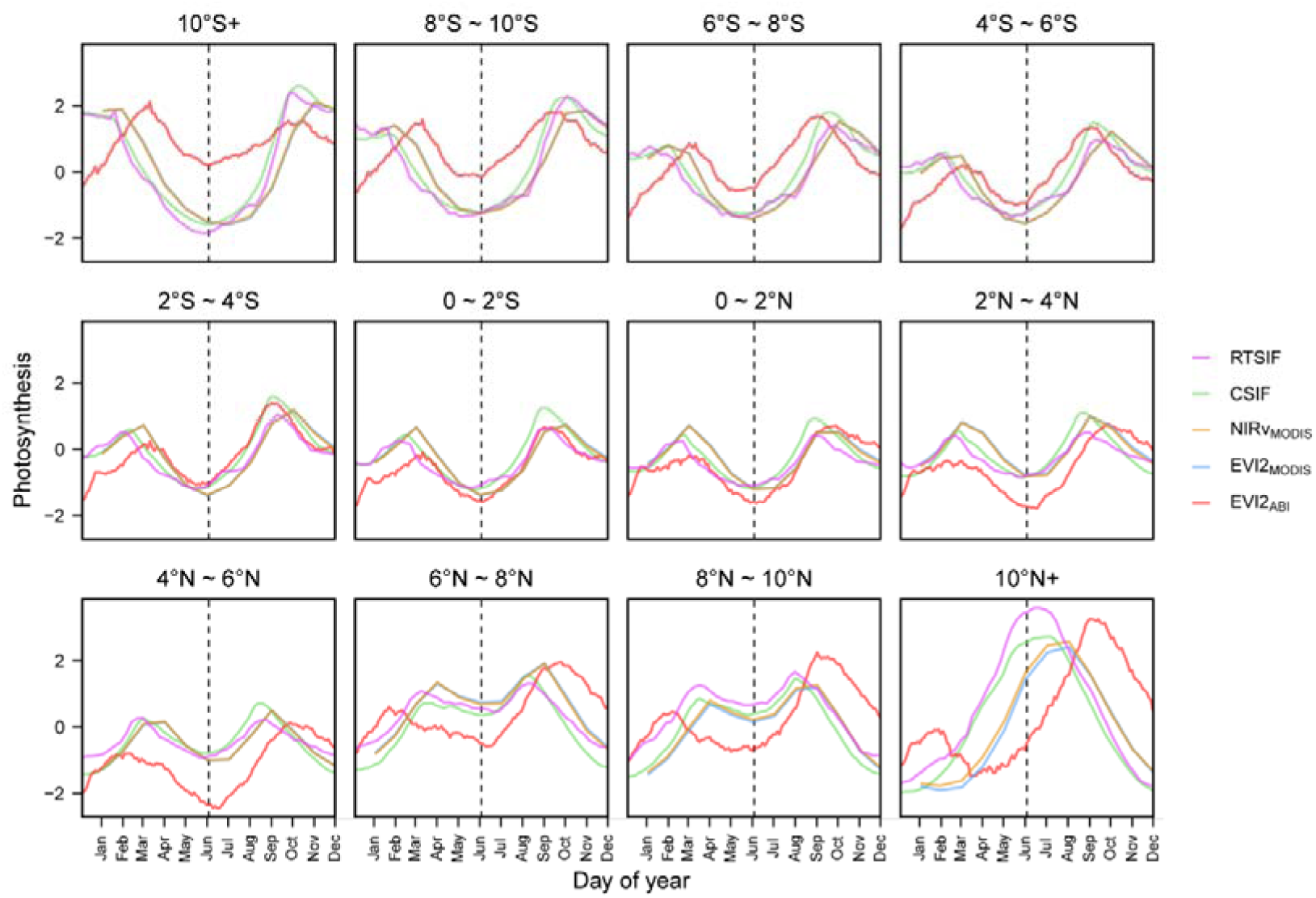
Photosynthetic seasonality along latitudinal gradients. For each dataset, values are obtained by subtracting the mean from the original values and then dividing by the standard deviation. Vertical dashed lines mark the half-year boundary (the 183rd day of the year).

**Fig. S3.**
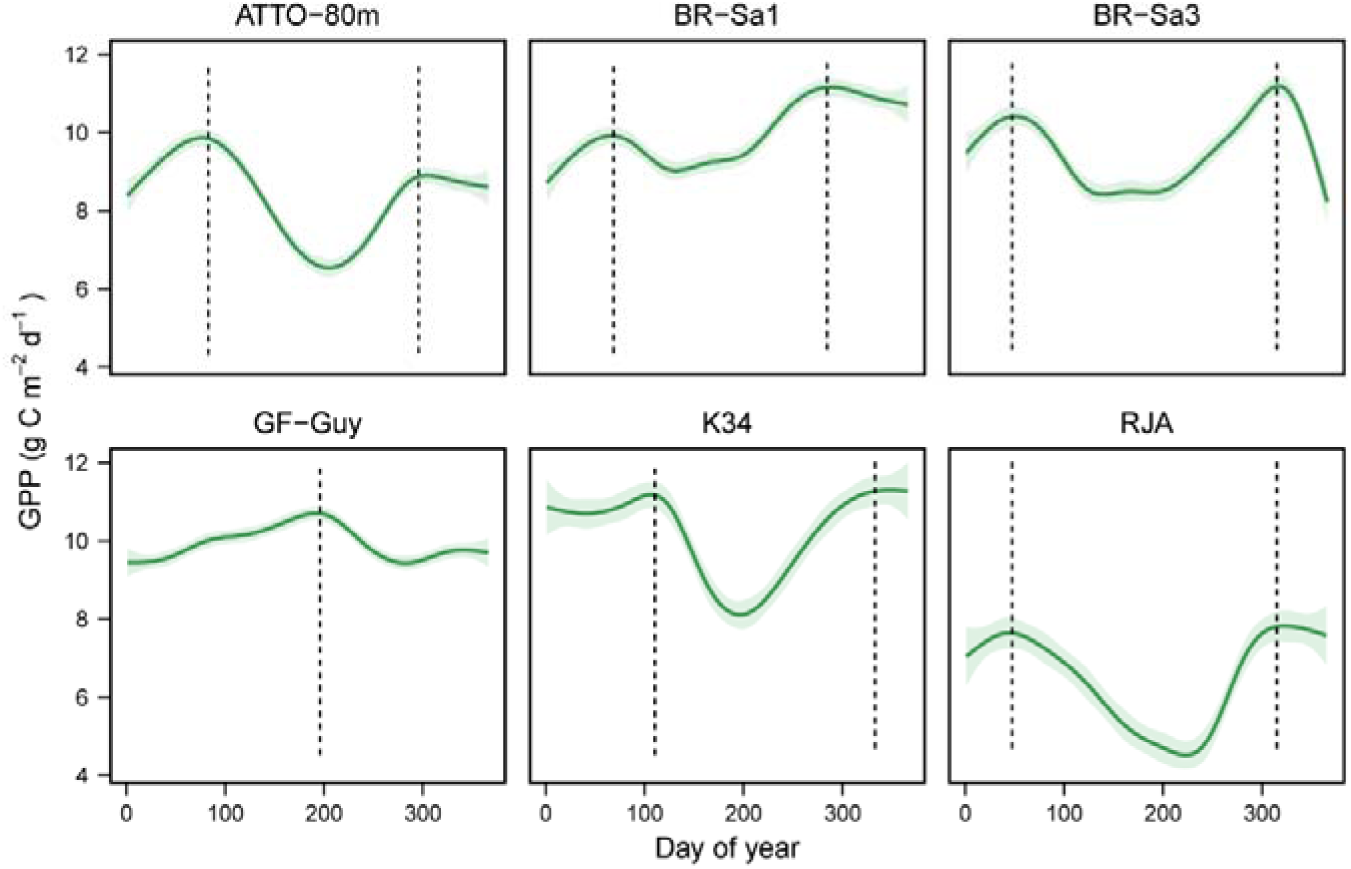
Photosynthetic seasonality at flux tower sites. Generalized additive models were used to fit the curves, with grey shading indicating 95% confidence intervals. Vertical dashed lines mark the timing of gross primary productivity (GPP) peaks.

**Fig. S4.**
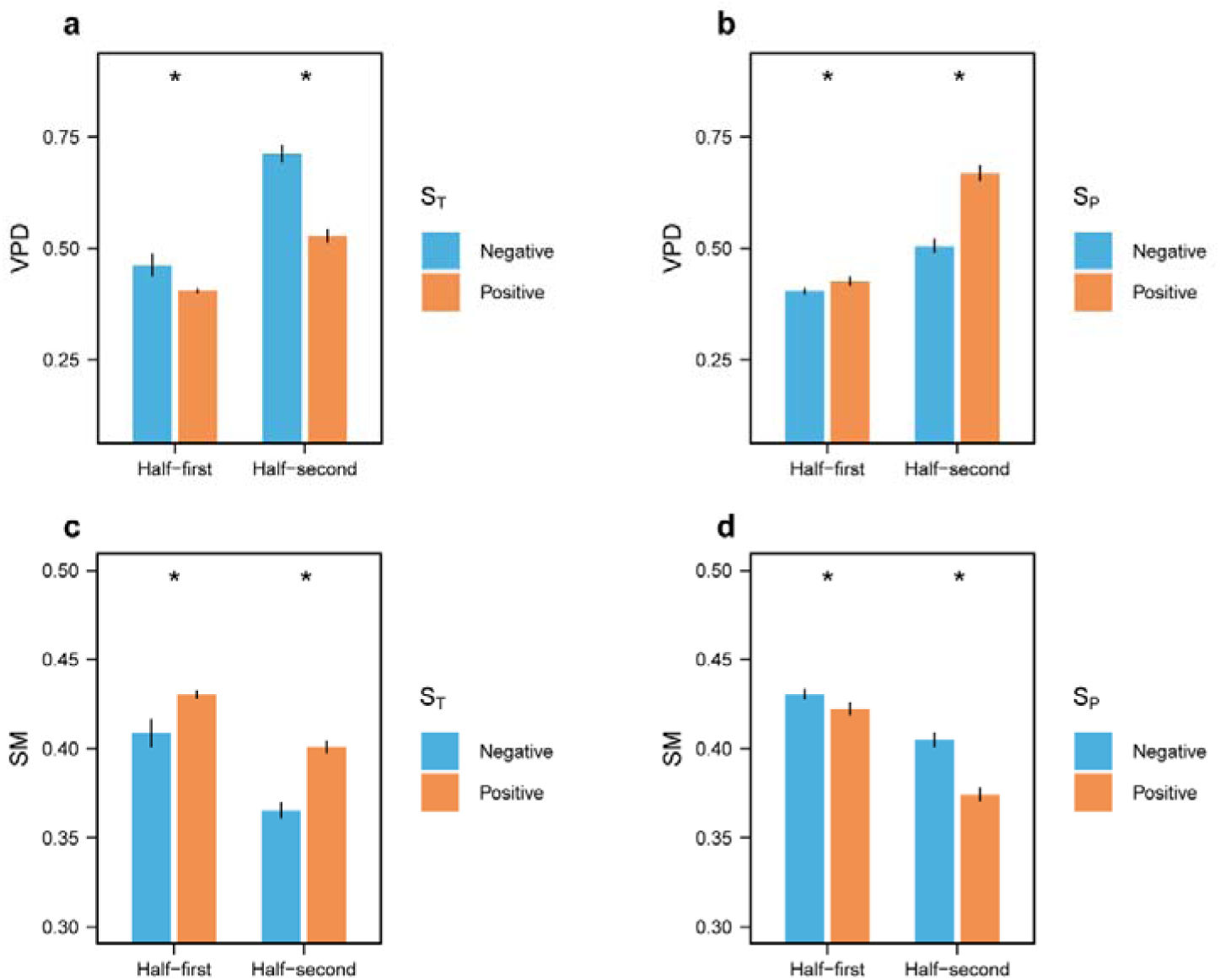
Comparison of VPD and SM between positive and negative hydrothermal sensitivities. The bars represent the mean values of VPD and SM, with error bars indicating the mean ± 5 × standard error (s.e.). The asterisks indicate statistically significant differences (Student’s *t*-test; *p* < 0.05).

**Fig. S5.**
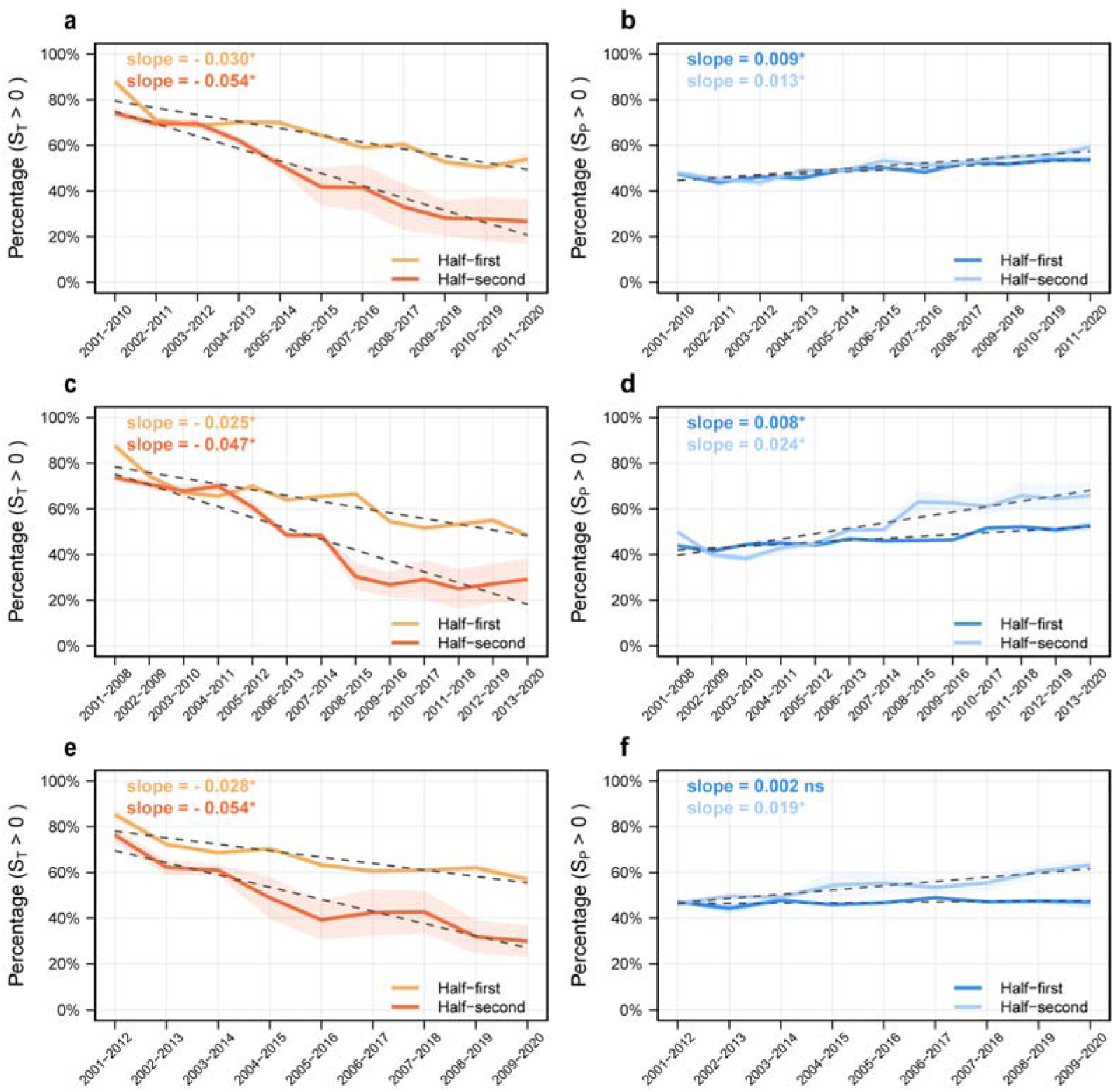
Temporal changes in hydrothermal sensitivities. a, b, Proportions of positive S_T_ and S_P_ obtained from ridge regression over time. **c-f**, Proportions of positive S_T_ and S_P_ obtained from 8-year (**c, d**) and 12-year (**e, f**) moving windows over time. The solid lines indicate the multi-dataset average, with the shaded area representing one standard deviation on either side of the mean. The dashed lines represent the linear regression fits, and the asterisks represent the slopes that are statistically significant (Student’s *t*-test; *p* < 0.05).

**Fig. S6.**
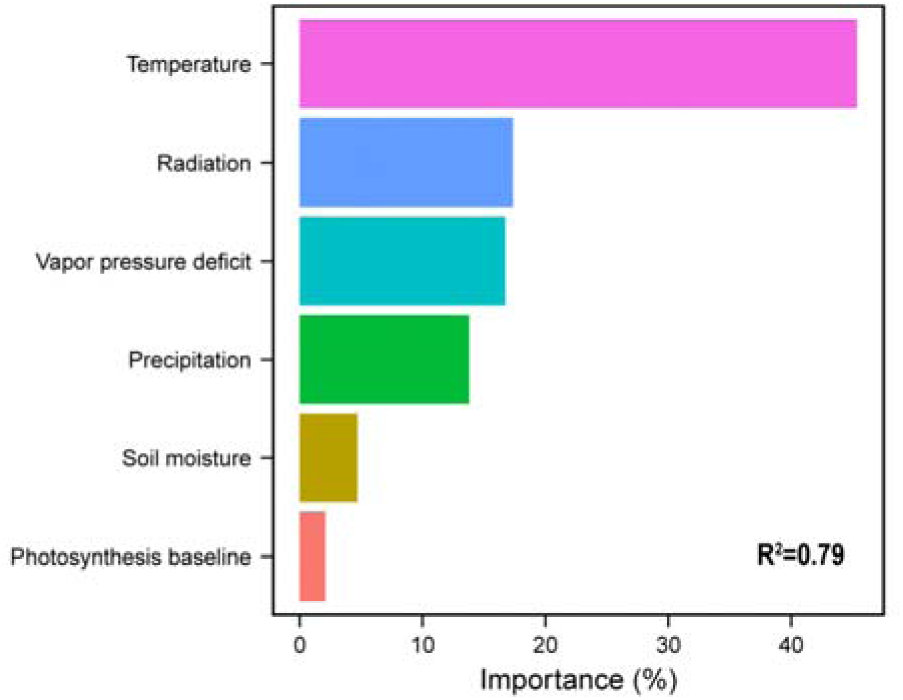
Importance of drivers influencing changes in T_opt_. The results are derived from a random forest model which explains 79% of the variation in T_opt_.

**Fig. S7.**
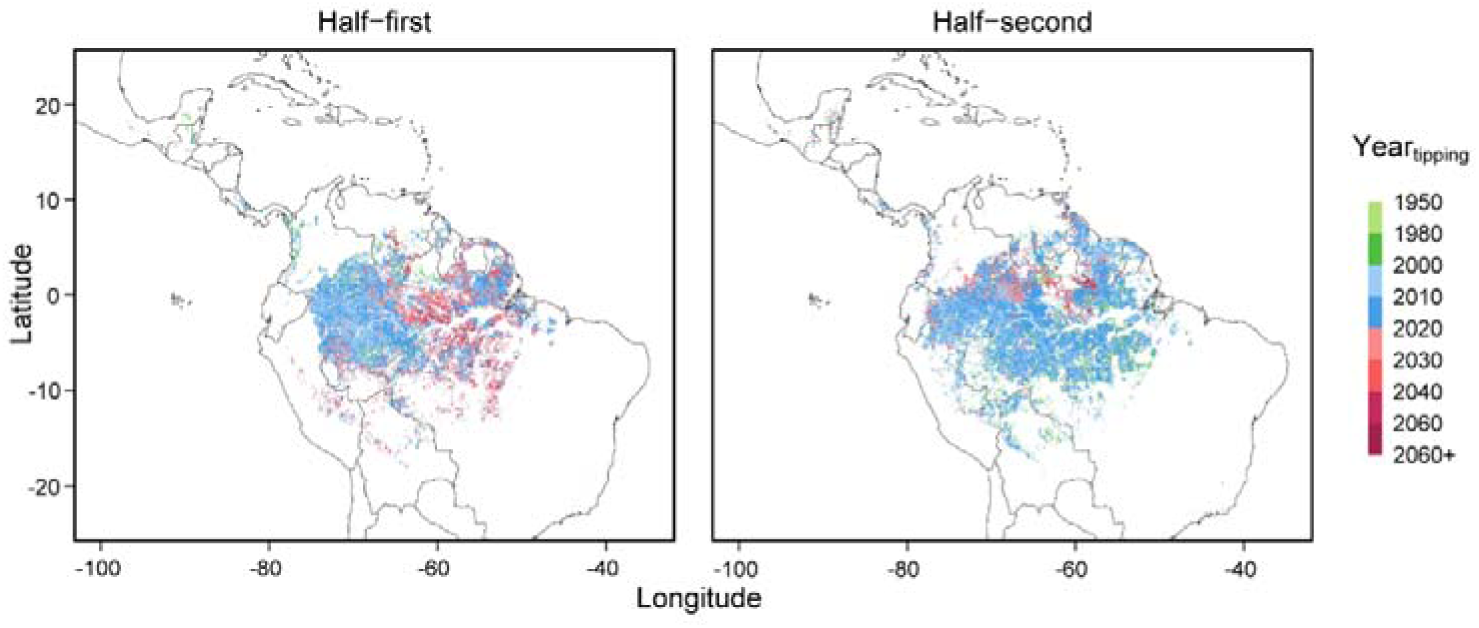
Spatial pattern of Year_tipping_. Year_tipping_ post-2020 here is derived under the SSP245 scenario.

**Fig. S8.**
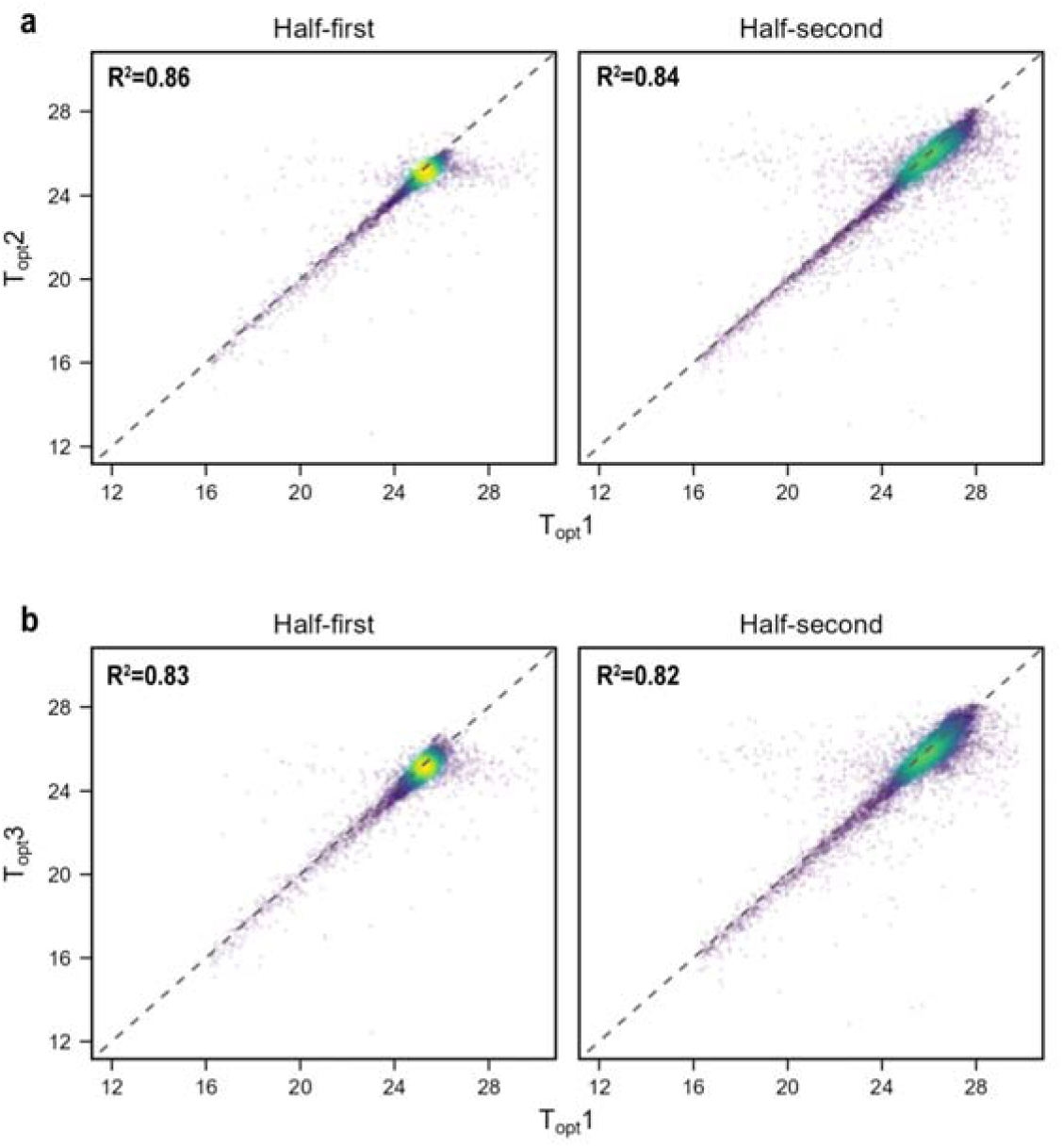
Comparison of T_opt_ obtained from different methods. T_opt_1 is calculated using quadratic functions, T_opt_2 is determined by Method2, and T_opt_3 is determined by Method3 (See Methods). The dashed 1:1 lines indicate perfect agreement between different T_opt_ estimates.

**Table S1.** Basic information about the flux towers used in this study.

| Site ID | Longitude | Latitude | Mean temperature (°C)<br>(First/second half) | Mean precipitation (mm)<br>(First/second half) | Available years |
| --- | --- | --- | --- | --- | --- |
| K34 | -60.20910 | -2.60900 | 25.16/26.33 | 1647.28 /788.82 | 2004, 2005, 2007, 2009,<br>2010, 2011 |
| RJA | -61.9331 | -10.078 | 24.70/25.81 | 1065.83/751.84 | 2001 |
| BR-Sa1 | -54.95889 | -2.85667 | 25.22/26.57 | 1749.14/936.65 | 2002-2006, 2008-2011 |
| BR-Sa3 | -54.97144 | -3.01803 | 25.02/26.60 | 1074.66/376.35 | 2001-2004 |
| GF-Guy | -52.92486 | 5.27877 | 25.30/25.92 | 2328.58/781.92 | 2004-2014 |
| ATTO-80m | -58.99 | -2.1441 | 25.60/27.17 | 1441.00/596.28 | 2014-2020 |

**Table S2.**
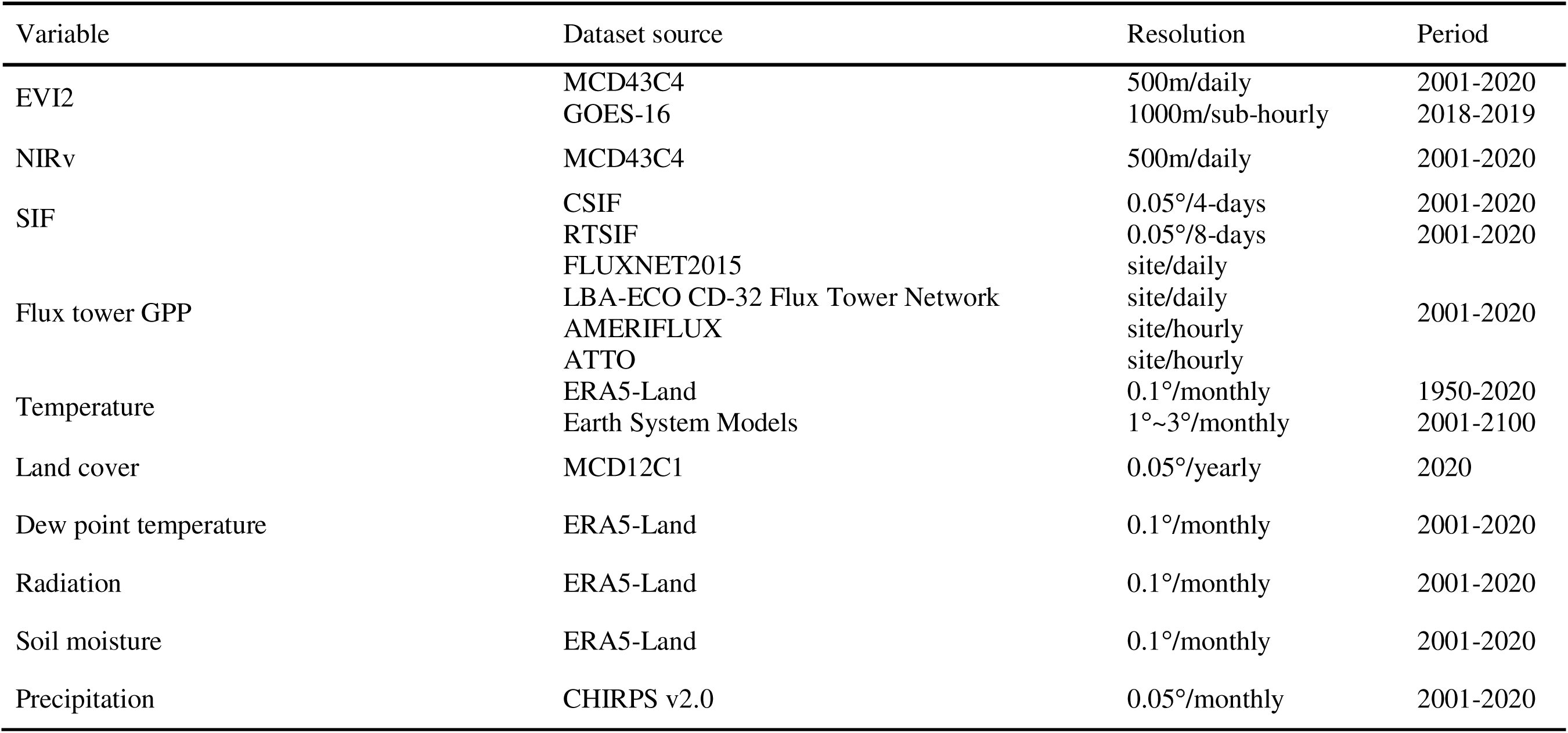
List of detailed information of the datasets used in this study.

## References

1. Saatchi S. S., Harris N. L., Brown S., Lefsky M., Mitchard E. T. et al. Benchmark map of forest carbon stocks in tropical regions across three continents. Proceedings of the national academy of sciences. 108, 9899–9904 (2011).

2. Nobre, C., A. Encalada, E. Anderson, F. Alcazar, et al. Amazon Assessment Report 2021 (2021).

3. Cox P. M., Betts R. A., Jones C. D., Spall S. A. & Totterdell I. J. Acceleration of global warming due to carbon-cycle feedbacks in a coupled climate model. Nature. 408, 184–187 (2000).

4. Marca□Zevallos M. J., Moulatlet G. M., Sousa T. R., Schietti J., Coelho L. d. S., et al. Local hydrological conditions influence tree diversity and composition across the Amazon basin. Ecography. 2022, e06125 (2022).

5. Mc□owell N., Allen C. D., Anderson□Teixeira K., Brando P., Brienen R. et al. Drivers and mechanisms of tree mortality in moist tropical forests. New Phytologist. 219, 851–869 (2018).

6. Park Williams A., Allen C. D., Macalady A. K., Griffin D., Woodhouse C. A. et al. Temperature as a potent driver of regional forest drought stress and tree mortality. Nature climate change. 3, 292–297 (2013).

7. Phillips O. L., Aragão L. E., Lewis S. L., Fisher J. B., Lloyd J. et al. Drought sensitivity of the Amazon rainforest. Science. 323, 1344–1347 (2009).

8. Reis S. M., Marimon B. S., Esquivel□Muelbert A., Marimon Jr B. H., Morandi P. S. et al. Climate and crown damage drive tree mortality in southern Amazonian edge forests. Journal of Ecology. 110, 876–888 (2022).

9. Gatti L. V., Basso L. S., Miller J. B., Gloor M., Gatti Domingues L. et al. Amazonia as a carbon source linked to deforestation and climate change. Nature. 595, 388–393 (2021).

10. Baccini A., Walker W., Carvalho L., Farina M., Sulla-Menashe D. et al. Tropical forests are a net carbon source based on aboveground measurements of gain and loss. Science. 358, 230–234 (2017).

11. Brienen R. J., Phillips O. L., Feldpausch T. R., Gloor E., Baker T. R. et al. Long-term decline of the Amazon carbon sink. Nature. 519, 344–348 (2015).

12. Hubau W., Lewis S. L., Phillips O. L., Affum-Baffoe K., Beeckman H. et al. Asynchronous carbon sink saturation in African and Amazonian tropical forests. Nature. 579, 80–87 (2020).

13. Phillips O. L., Brienen R. J. & collaboration R. Carbon uptake by mature Amazon forests has mitigated Amazon nations’ carbon emissions. Carbon Balance Manag. 12, 1–9 (2017).

14. Li Q., Chen X., Yuan W., Lu H., Shen R. et al. Remote sensing of seasonal climatic constraints on leaf phenology across pantropical evergreen forest biome. Earth’s Future. 9, e2021EF002160 (2021).

15. Yang X., Wu J., Chen X., Ciais P., Maignan F. et al. A comprehensive framework for seasonal controls of leaf abscission and productivity in evergreen broadleaved tropical and subtropical forests. The Innovation. 2, 100154 (2021).

16. Chen R., Liu L. & Liu X. The negative impact of excessive moisture contributes to the seasonal dynamics of photosynthesis in Amazon moist forests. Earth’s Future. 10, e2021EF002306 (2022).

17. Chen X., Maignan F., Viovy N., Bastos A., Goll D. et al. Novel representation of leaf phenology improves simulation of Amazonian evergreen forest photosynthesis in a land surface model. Journal of Advances in Modeling Earth Systems. 12, e2018MS001565 (2020).

18. Zhang X., Shen Y., Gao S., Wang W. & Schaaf C. Diverse Responses of Multiple Satellite□Derived Vegetation Greenup Onsets to Dry Periods in the Amazon. Geophysical Research Letters. 49, e2022GL098662 (2022).

19. Girardin C. A., Malhi Y., Doughty C. E., Metcalfe D. B., Meir P. et al. Seasonal trends of Amazonian rainforest phenology, net primary productivity, and carbon allocation. Global Biogeochemical Cycles. 30, 700–715 (2016).

20. Wright S. J. & van Schaik C. P. Light and the phenology of tropical trees. The American Naturalist. 143, 192–199 (1994).

21. Shaw P. Rainfall, leafing phenology and sunrise time as potential Zeitgeber for the bimodal, dry season laying pattern of an African rain forest tit (Parus fasciiventer). J Ornithol. 158, 263–275 (2017).

22. Yan H., Wang S. Q., da Rocha H. R., Rap A., Bonal D. et al. Simulation of the unexpected photosynthetic seasonality in Amazonian evergreen forests by using an improved diffuse fraction□based light use efficiency model. Journal of Geophysical Research: Biogeosciences. 122, 3014–3030 (2017).

23. Liu L., Ciais P., Maignan F., Zhang Y., Viovy N. et al. Solar radiation triggers the bimodal leaf phenology of central African evergreen broadleaved forests. Journal of Advances in Modeling Earth Systems. 16, e2023MS004014 (2024).

24. Terasaki Hart D. E., Bùi T.-N., Di Maggio L. & Wang I. J. Global phenology maps reveal the drivers and effects of seasonal asynchrony. Nature. 1–8 (2025).

25. Kattge J. & Knorr W. Temperature acclimation in a biochemical model of photosynthesis: a reanalysis of data from 36 species. Plant, cell & environment. 30, 1176–1190 (2007).

26. Lloyd J. & Farquhar G. D. Effects of rising temperatures and [CO_2_] on the physiology of tropical forest trees. Philosophical Transactions of the Royal Society B: Biological Sciences. 363, 1811–1817 (2008).

27. Doughty C. E., Keany J. M., Wiebe B. C., Rey-Sanchez C., Carter K. R. et al. Tropical forests are approaching critical temperature thresholds. Nature. 621, 105–111 (2023).

28. Mau A. C., Reed S. C., Wood T. E. & Cavaleri M. A. Temperate and tropical forest canopies are already functioning beyond their thermal thresholds for photosynthesis. Forests. 9, 47 (2018).

29. Doughty C. E. & Goulden M. L. Are tropical forests near a high temperature threshold? Journal of Geophysical Research: Biogeosciences. 113, G00B07 (2008).

30. Doughty R., Köhler P., Frankenberg C., Magney T. S., Xiao X. et al. TROPOMI reveals dry-season increase of solar-induced chlorophyll fluorescence in the Amazon forest. Proceedings of the National Academy of Sciences. 116, 22393–22398 (2019).

31. Hilker T., Lyapustin A. I., Tucker C. J., Hall F. G., Myneni R. B. et al. Vegetation dynamics and rainfall sensitivity of the Amazon. Proceedings of the National Academy of Sciences. 111, 16041–16046 (2014).

32. Huete A. R., Didan K., Shimabukuro Y. E., Ratana P., Saleska S. R. et al. Amazon rainforests green□up with sunlight in dry season. Geophysical research letters. 33, L06405 (2006).

33. Morton D. C., Nagol J., Carabajal C. C., Rosette J., Palace M. et al. Amazon forests maintain consistent canopy structure and greenness during the dry season. Nature. 506, 221–224 (2014).

34. Samanta A., Ganguly S. & Myneni R. B. MODIS enhanced vegetation index data do not show greening of Amazon forests during the 2005 drought. The New Phytologist. 189, 11–15 (2011).

35. Signori-Müller C., Oliveira R. S., Barros F. d. V., Tavares J. V., Gilpin M., et al. Non-structural carbohydrates mediate seasonal water stress across Amazon forests. Nature Communications. 12, 2310 (2021).

36. Flores B. M., Montoya E., Sakschewski B., Nascimento N., Staal A. et al. Critical transitions in the Amazon forest system. Nature. 626, 555–564 (2024).

37. Zemp D. C., Schleussner C.-F., Barbosa H. M., Hirota M., Montade V. et al. Self-amplified Amazon forest loss due to vegetation-atmosphere feedbacks. Nature communications. 8, 14681 (2017).

38. Chai Y., Martins G., Nobre C., von Randow C., Chen T. et al. Constraining Amazonian land surface temperature sensitivity to precipitation and the probability of forest dieback. NPJ Climate and Atmospheric Science. 4, 6 (2021).

39. Fu R., Yin L., Li W., Arias P. A., Dickinson R. E. et al. Increased dry-season length over southern Amazonia in recent decades and its implication for future climate projection. Proceedings of the National Academy of Sciences. 110, 18110–18115 (2013).

40. Xu H., Lian X., Slette I. J., Yang H., Zhang Y. et al. Rising ecosystem water demand exacerbates the lengthening of tropical dry seasons. Nature Communications. 13, 4093 (2022).

41. Guo J., Hu S. & Guan Y. Regime shifts of the wet and dry seasons in the tropics under global warming. Environmental Research Letters. 17, 104028 (2022).

42. Valeriano C., Gutiérrez E., Colangelo M., Gazol A., Sánchez-Salguero R. et al. Seasonal precipitation and continentality drive bimodal growth in Mediterranean forests. Dendrochronologia. 78, 126057 (2023).

43. Tumajer J., Shishov V. V., Ilyin V. A. & Camarero J. J. Intra-annual growth dynamics of Mediterranean pines and junipers determines their climatic adaptability. Agricultural and Forest Meteorology. 311, 108685 (2021).

44. Campelo F., Ribas M. & Gutiérrez E. Plastic bimodal growth in a Mediterranean mixed-forest of Quercus ilex and Pinus halepensis. Dendrochronologia. 67, 125836 (2021).

45. Tercek M. T., Gross J. E. & Thoma D. P. Robust projections and consequences of an expanding bimodal growing season in the western United States. Ecosphere. 14, e4530 (2023).

46. Weiss J. L., Gutzler D. S., Coonrod J. E. A. & Dahm C. N. Long-term vegetation monitoring with NDVI in a diverse semi-arid setting, central New Mexico, USA. Journal of Arid Environments. 58, 249–272 (2004).

47. Notaro M., Liu Z., Gallimore R. G., Williams J. W., Gutzler D. S. et al. Complex seasonal cycle of ecohydrology in the Southwest United States. Journal of Geophysical Research: Biogeosciences. 115, G4 (2010).

48. Grossiord C., Buckley T. N., Cernusak L. A., Novick K. A., Poulter B. et al. Plant responses to rising vapor pressure deficit. New phytologist. 226, 1550–1566 (2020).

49. Slot M., Rifai S. W., Eze C. E. & Winter K. The stomatal response to vapor pressure deficit drives the apparent temperature response of photosynthesis in tropical forests. New Phytologist. 244, 1238–1249 (2024).

50. Wang R., Li L., Gentine P., Zhang Y., Chen J. et al. Recent increase in the observation-derived land evapotranspiration due to global warming. Environmental Research Letters. 17, 024020 (2022).

51. Sperry J. S. & Tyree M. T. Mechanism of water stress-induced xylem embolism. Plant Physiol. 88, 581–587 (1988).

52. Tyree M. T. & Dixon M. A. Water stress induced cavitation and embolism in some woody plants. Physiologia Plantarum. 66, 397–405 (1986).

53. Choat B., Jansen S., Brodribb T. J., Cochard H., Delzon S. et al. Global convergence in the vulnerability of forests to drought. Nature. 491, 752–755 (2012).

54. Sharkey T. D. Effects of moderate heat stress on photosynthesis: importance of thylakoid reactions, rubisco deactivation, reactive oxygen species, and thermotolerance provided by isoprene. Plant, cell & environment. 28, 269–277 (2005).

55. Lewis S. L., Brando P. M., Phillips O. L., Van Der Heijden G. M. & Nepstad D. The 2010 amazon drought. Science. 331, 554-554 (2011).

56. Zeng N., Yoon J.-H., Marengo J. A., Subramaniam A., Nobre C. A. et al. Causes and impacts of the 2005 Amazon drought. Environmental Research Letters. 3, 014002 (2008).

57. Song F., Dong H., Wu L., Leung L. R., Lu J. et al. Hot season gets hotter due to rainfall delay over tropical land in a warming climate. Nature Communications. 16, 2188 (2025).

58. Staal A., Tuinenburg O. A., Bosmans J. H., Holmgren M., van Nes E. H. et al. Forest-rainfall cascades buffer against drought across the Amazon. Nature Climate Change. 8, 539–543 (2018).

59. Cui J., Lian X., Huntingford C., Gimeno L., Wang T. et al. Global water availability boosted by vegetation-driven changes in atmospheric moisture transport. Nature Geoscience. 15, 982–988 (2022).

60. Leroi A. M., Bennett A. F. & Lenski R. E. Temperature acclimation and competitive fitness: an experimental test of the beneficial acclimation assumption. Proceedings of the National Academy of Sciences. 91, 1917–1921 (1994).

61. Yamori W., Hikosaka K. & Way D. A. Temperature response of photosynthesis in C_3_, C_4_, and CAM plants: temperature acclimation and temperature adaptation. Photosynthesis research. 119, 101–117 (2014).

62. Huang M., Piao S., Ciais P., Peñuelas J., Wang X. et al. Air temperature optima of vegetation productivity across global biomes. Nature ecology & evolution. 3, 772–779 (2019).

63. Zhang Y., Piao S., Sun Y., Rogers B. M., Li X. et al. Future reversal of warming-enhanced vegetation productivity in the Northern Hemisphere. Nature Climate Change. 12, 581–586 (2022).

64. Zhai L., Will R. E. & Zhang B. Structural diversity is better associated with forest productivity than species or functional diversity. Ecology. 105, e4269 (2024).

65. Liu X., Feng Y., Hu T., Luo Y., Zhao X. et al. Enhancing ecosystem productivity and stability with increasing canopy structural complexity in global forests. Science Advances. 10, eadl1947 (2024).

66. Gu H., Qiao Y., Xi Z., Rossi S., Smith N. G. et al. Warming-induced increase in carbon uptake is linked to earlier spring phenology in temperate and boreal forests. Nature Communications. 13, 3698 (2022).

67. Springer C. J. & Ward J. K. Flowering time and elevated atmospheric CO2. New Phytologist. 176, 243–255 (2007).

68. Greer D. H., Cirillo C. & Norling C. L. Temperature-dependence of carbon acquisition and demand in relation to shoot and fruit growth of fruiting kiwifruit (Actinidia deliciosa) vines grown in controlled environments. Funct. Plant Biol. 30, 927–937 (2003).

69. Aragão L. E., Anderson L. O., Fonseca M. G., Rosan T. M., Vedovato L. B. et al. 21st Century drought-related fires counteract the decline of Amazon deforestation carbon emissions. Nature communications. 9, 536 (2018).

70. Zhang Z., Cescatti A., Wang Y.-P., Gentine P., Xiao J. et al. Large diurnal compensatory effects mitigate the response of Amazonian forests to atmospheric warming and drying. Science advances. 9, eabq4974 (2023).

71. Green J., Berry J., Ciais P., Zhang Y. & Gentine P. Amazon rainforest photosynthesis increases in response to atmospheric dryness. Science Advances. 6, eabb7232 (2020).

72. Chen S., Stark S. C., Nobre A. D., Cuartas L. A., de Jesus Amore D. et al. Amazon forest biogeography predicts resilience and vulnerability to drought. Nature. 631, 111–117 (2024).

73. Chen X., Huang Y., Nie C., Zhang S., Wang G. et al. A long-term reconstructed TROPOMI solar-induced fluorescence dataset using machine learning algorithms. Scientific Data. 9, 427 (2022).

74. Zhang Y., Joiner J., Alemohammad S. H., Zhou S. & Gentine P. A global spatially contiguous solar-induced fluorescence (CSIF) dataset using neural networks. Biogeosciences. 15, 5779–5800 (2018).

75. Badgley G., Field C. B. & Berry J. A. Canopy near-infrared reflectance and terrestrial photosynthesis. Science advances. 3, e1602244 (2017).

76. Badgley G., Anderegg L. D., Berry J. A. & Field C. B. Terrestrial gross primary production: Using NIRV to scale from site to globe. Global change biology. 25, 3731–3740 (2019).

77. Jiang Z., Huete A. R., Didan K. & Miura T. Development of a two-band enhanced vegetation index without a blue band. Remote sensing of Environment. 112, 3833–3845 (2008).

78. Román M. O., Justice C., Paynter I., Boucher P. B., Devadiga S. et al. Continuity between NASA MODIS Collection 6.1 and VIIRS Collection 2 land products. Remote Sensing of Environment. 302, 113963 (2024).

79. Campagnolo M. L., Sun Q., Liu Y., Schaaf C., Wang Z. et al. Estimating the effective spatial resolution of the operational BRDF, albedo, and nadir reflectance products from MODIS and VIIRS. Remote Sensing of Environment. 175, 52–64 (2016).

80. Roy D. P., Li Z., Giglio L., Boschetti L. & Huang H. Spectral and diurnal temporal suitability of GOES Advanced Baseline Imager (ABI) reflectance for burned area mapping. IJAEO. 96, 102271 (2021).

81. Hashimoto H., Wang W., Dungan J. L., Li S., Michaelis A. R. et al. New generation geostationary satellite observations support seasonality in greenness of the Amazon evergreen forests. Nature Communications. 12, 684 (2021).

82. Schaaf C. B., Gao F., Strahler A. H., Lucht W., Li X. et al. First operational BRDF, albedo nadir reflectance products from MODIS. Remote sensing of Environment. 83, 135–148 (2002).

83. Roy D. P., Lewis P., Schaaf C., Devadiga S. & Boschetti L. The global impact of clouds on the production of MODIS bidirectional reflectance model-based composites for terrestrial monitoring. IEEE Geoscience and Remote Sensing Letters. 3, 452–456 (2006).

84. Shen Y., Zhang X., Gao S., Zhang H. K., Schaaf C. et al. Analyzing GOES-R ABI BRDF-adjusted EVI2 time series by comparing with VIIRS observations over the CONUS. Remote Sensing of Environment. 302, 113972 (2024).

85. Andreae M. O., Acevedo O. C., Araùjo A., Artaxo P., Barbosa C. G. et al. The Amazon Tall Tower Observatory (ATTO): overview of pilot measurements on ecosystem ecology, meteorology, trace gases, and aerosols. Atmospheric Chemistry and Physics. 15, 10723–10776 (2015).

86. Murray F. W. On the computation of saturation vapor pressure. Journal of Applied Meteorology and Climatology. 6, 203–204 (1967).

87. Gray J., Sulla-Menashe D. & Friedl M. A. User guide to collection 6 modis land cover dynamics (mcd12q2) product. *NASA EOSDIS Land Processes DAAC: Missoula, MT*, USA. 6, 1–8 (2019).

88. Hoerl A. E. & Kennard R. W. Ridge regression: biased problems nonorthogonal estimation for nonorthogonal problems. Technometrics. 42, 80–86 (2000).

89. Sendall K. M., Reich P. B., Zhao C., Jihua H., Wei X. et al. Acclimation of photosynthetic temperature optima of temperate and boreal tree species in response to experimental forest warming. Global Change Biology. 21, 1342–1357 (2015).

90. Battaglia M., Beadle C. & Loughhead S. Photosynthetic temperature responses of Eucalyptus globulus and Eucalyptus nitens. Tree physiology. 16, 81–89 (1996).

91. Li X., Huntingford C., Wang K., Cui J., Xu H. et al. Increased crossing of thermal stress thresholds of vegetation under global warming. Global Change Biology. 30, e17406 (2024).

